# Mitotic chromosome condensation resets chromatin to maintain transcriptional homeostasis

**DOI:** 10.1101/2022.05.11.491439

**Authors:** Lucía Ramos-Alonso, Petter Holland, Stéphanie Le Gras, Xu Zhao, Bernard Jost, Magnar Bjørås, Yves Barral, Jorrit M. Enserink, Pierre Chymkowitch

## Abstract

Mitotic entry correlates with the condensation of the chromosomes, remodeling of histone modifications, exclusion of transcription factors from DNA and the broad downregulation of transcription. However, whether mitotic condensation influences transcription in the subsequent interphase is unknown. Here, we show that preventing one chromosome to condense during mitosis causes it to fail resetting transcription. Rather it diverted the transcription machinery and underwent unscheduled initiation of gene expression. This caused the activation of inducible transcriptional programs, such as the *GAL* genes, even in absence of the relevant stimuli. Strikingly, aberrant gene expression persisted into the next interphase. Thus, our study identifies the maintenance of transcriptional homeostasis as an unexpected and yet unexplored function of mitotic chromosome condensation.

**One-Sentence Summary:** Mitotic chromatin condensation resets the transcriptome to protect cells from transcriptional drifting after anaphase.

## Main Text

During mitotic entry, eukaryotic chromosomes undergo a drastic compaction process, called chromosome condensation, that makes them manageable units for the mitotic spindle to segregate them between the two daughter cells (*1*). In addition, mitotic chromosome compaction remodels the genome architecture of interphase, preventing the association of most transcription- and chromatin regulators to *cis* regulatory elements (*2, 3*). This correlates with a near-complete shutdown of the expression of interphase genes, except for certain housekeeping genes, which remain weakly transcribed to maintain basic cellular functions (*3, 4*). At the end of mitosis, interphase gene expression programs are reestablished in a precise and timely coordinated manner. In a first wave, cells enhance primarily the expression of housekeeping genes, whereas genes involved in the maintenance of cellular identity and functions are reactivated in a second wave (*4, 5*). This correlates with the timely reestablishment of interphase chromatin architecture and reloading of the transcription machinery. Although mitotic promoter bookmarking and histone modifications participate, the mechanisms enforcing controlled genome reactivation at mitotic exit remain unclear (*5–7*). While the dramatic transcriptional silencing of mitotic chromosomes has been observed and accepted a long time ago, it is typically regarded as a passive consequence of mitotic chromatin condensation and it remains unclear whether it has any biological significance beyond M phase.

We recently reported that the excision of a single yeast centromere, beyond preventing kinetochore assembly, prevented the condensation of the entire chromosome (*8, 9*). This study established that at least in budding yeast condensation takes place in a chromosome-autonomous manner, is initiated at centromeres, and proceeds through the sequential mobilization of the kinase Ipl1, the protein shugoshin (Sgo1) and the sirtuin Hst2. In turn these proteins mediated histone deacetylation, chromatin compaction and condensin-dependent contraction of the chromosome arms (*8–10*). Here, we used the same approach, namely the controlled excision of the centromere of yeast chromosome IV (*CEN4*), to investigate the consequence of preventing condensation of a single chromosome on transcription in the subsequent cell cycle. The centromere of chromosome IV flanked with lox recombination sites (*CEN4*\*; Fig. 1A) (*9, 11*), was excised on demand through activation of a chimeric Cre protein fused to an estradiol-binding domain (Cre-EBD). Concretely, *CEN4\** was quickly and efficiently excised from the chromosome in a majority of the cells 30 minutes after estradiol treatment (*12*) (fig. S1A).

**Fig. 1.**
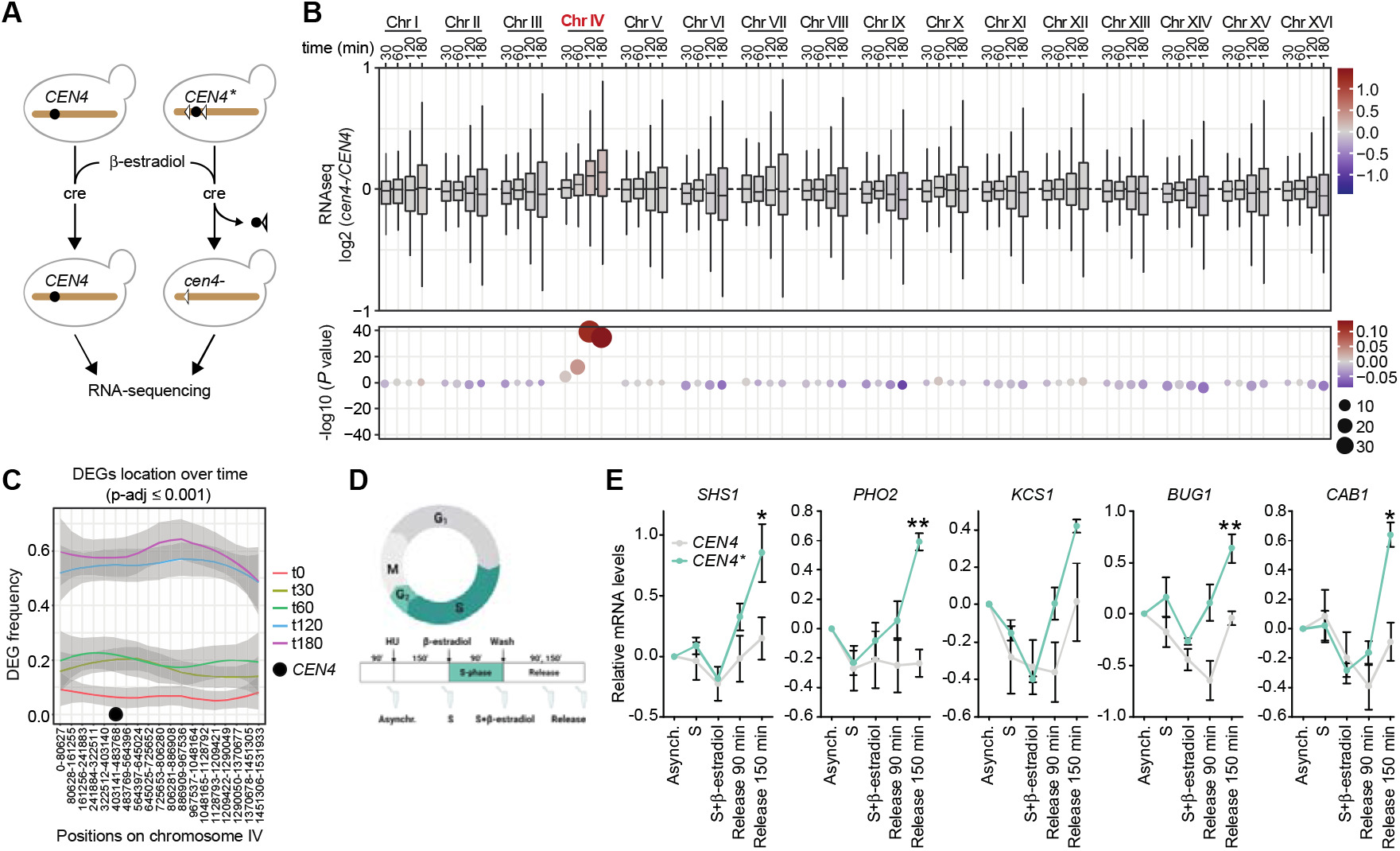
Failure to condense chromatin at mitotic entry results in general upregulation of gene expression during interphase. (**A**) The centromere excision assay. (**B**) RNAseq assessment of gene expression at each chromosome and at different time points after *CEN4\** excision. (**C**) Frequency of DEGs (*P*-adj ≤ 0.001) at chromosome IV over time after *CEN4\** excision. (**D**) Layout of *CEN4\** excision in S-phase arrest and release experiments. (**E**) RT-qPCR analysis of the expression of five selected genes from chromosome IV at different time points before and after S-phase arrest and release *(see D). *P* values < 0.05; \*\**P* values < 0.01

RNA-sequencing (RNAseq) at several time points after *CEN4\** excision (see fig. S1B for PCA) enabled us to analyze the effect of *CEN4\** excision on gene expression on all yeast chromosomes, using increasingly stringent *P*-adj cutoffs. Strikingly, in addition to triggering global transcriptome deregulation, *CEN4\** excision was followed by a specific, significant and progressive upregulation of transcription of genes located on chromosome IV. The amplitude (Log2 FC) of this raise and its significance were much more pronounced than at any other chromosome (Fig. 1B, fig. S1, C to E, table S4 and table S5). The enrichment of upregulated genes at chromosome IV was validated by unbiased hierarchical clustering. These unbiassed analyses identified a large cluster of upregulated genes that was significantly enriched for genes located on chromosome IV (fig. S2, A to D). Other clusters contained genes that were either upregulated, downregulated or unaffected upon *CEN4\** excision and were not enriched for genes belonging to chromosome IV (fig. S2, A to D). Consistent with the notion that the localisation of these genes drove their upregulation, Gene Ontology (GO) analysis of the genes deregulated upon *CEN4\** excision could not classify the deregulated genes from chromosome IV into any specific pathway (except a few genes involved in “cellular response to DNA damage stimulus”, fig. S3, A and B).

To study this effect in more detail we focused on *GAL3*, which is located on chromosome IV. Its product Gal3p is a galactose-inducible transcription activator that promotes the expression of *GAL1, GAL2, GAL7* and *GAL10* (*13*) (fig. S4A). However, even in absence of galactose *GAL3* was strongly and increasingly activated over time upon *CEN4\** excision (fig. S4A). Interestingly, its target genes, which are located on chromosome II and XII, also became upregulated (fig. S4A), albeit less intensely and slower. A similar effect was observed for example for the *INO2* gene, which encodes an activator of phospholipid synthesis genes (*14*) (fig. S4B). Thus, loss of *CEN4\** unleashes the expression of chromosome IV genes, even in absence of cognate signals, occasionally resulting in the increased expression of downstream genes located on other chromosomes (fig. S4D).

Interestingly, in addition to causing aberrant expression of specific transcriptional programs, such as the *GAL* or *INO2* systems, excision of *CEN4\** affected several genes located outside chromosome IV that are involved in regulation of translation, ribosome biogenesis and metabolism. These pathways are often downregulated in response to stress, and partially overlapped with the environmental stress response program (ESR) (*15, 16*) (fig. S3, A and B and fig. S4C). We hypothesized that, similar to the *GAL* system, activation of the ESR was due to increased expression of ESR regulators located on chromosome IV. Although we cannot exclude this possibility, we note the main activators of ESR, *MSN2* and *MSN4*, are upregulated after *CEN4\** excision (Data S1), but are located outside chromosome IV. This suggests that ESR may be induced in response to centromere excision. Importantly, excision of a small patch of control DNA from one of the arms of chromosome IV using the same cre/lox system (fig. S5, A and B) had no effect on gene expression (fig. S5, C and D), demonstrating that transcriptional deregulation is specific to the excision of *CEN4\**. Taken together, these data show that centromeres suppress unscheduled activation of transcription *in cis*, and that loss of a single centromere induces the ESR.

Interestingly, gene deregulation at chromosome IV was partially dependent on the distance of genes from *CEN4\**. We did not observe any enrichment for any chromosomal region when we plotted all genes identified by RNAseq according to their position on chromosome IV (fig. S6A). Likewise, plotting the log2 FC values of the differentially expressed genes (*P* ≤ 0.001) against their position relative to *CEN4* revealed that the amplitude of gene expression changes (Log2 FC) is not linked to the location of genes on chromosome IV (see Pearson correlation score in fig. S6B). However, plotting the density of DEGs relative to the position of *CEN4\** over time revealed a time-dependent increase in the density of upregulated genes that peaked between 60 min and 120 min after *CEN4\** excision, and which appeared to propagate in wave-like patterns spreading from *CEN4* to the telomeres (Fig. 1C and ‘DEGs Up’ in fig. S6C). In contrast, the small group of downregulated genes at chromosome IV did not display a comparable increase over time, neither in density nor spreading (fig. S6C, ‘DEGs Down’). Thus, transcriptional upregulation in response to *CEN4\** excision appears to spread progressively from the centromeric region towards the end of the chromosome arms of chromosome IV.

To confirm that overexpression of genes on chromosome IV was the consequence of the mitotic condensation defect induced by *CEN4\** excision, we synchronized the cell cycle prior to chromosome condensation using hydroxyurea (HU), which arrests yeast cells in S phase right before anaphase onset (S phase partially overlaps with early phases of mitosis in budding yeast (*17–19*)). We then released the synchronized cells in presence or absence of β-estradiol, and analyzed the timing of upregulation of a panel of strongly upregulated genes (Fig. 1D). The expression of these genes was identical in asynchronous and in HU-arrested *CEN4* or *CEN4\** cells (Fig. 1E). Similar results were obtained with cells arrested in G1 using alpha-factor, or in prometaphase using nocodazole (fig. S7, A to D). In contrast, transcriptional activation in response to *CEN4\** excision was strong in cells that had been released from HU arrest for 90 min and 150 min and that had completed mitosis (Fig. 1E and fig. S7, E and F). Thus, progression through mitosis is required for centromere excision to cause aberrant gene upregulation.

We previously showed that the histone deacetylase Hst2 operates downstream of centromeres in chromatin condensation during mitotic entry (*8, 9*). Consistent with this and with overexpression of chromosome IV being due to a condensation defect, the occurrence of H4K12ac and H4K16ac substantially increased over time after *CEN4\** excision on chromosome IV, specifically, as determined by ChIP-sequencing (ChIPseq) (see fig. S8, A and B for PCA). Importantly, the other chromosomes remained largely unaffected (Fig. 2, A to D, fig. S9A and B and fig. S10, A and B). Histone acetylation started increasing near *CEN4\** before spreading to each chromosome arm over time (fig. S12A), similar to the gene expression patterns described above. Crucially, ATACseq experiments showed that increased histone acetylation at chromosome IV correlated with a strong increase in chromatin accessibility (Fig. 2E and fig. S10C). Thus, centromere inactivation causes aberrantly high levels of histones acetylation and chromatin opening on the affected chromosome, increasing the accessibility of DNA to the transcription machinery. Supporting this interpretation, the occurrence of the active promoter mark H3K4me3 was significantly increased at promoters on chromosome IV after β-estradiol treatment, specifically (Fig. 3, A and B, fig. S9, A and B, fig. S11A and fig. S8C for PCA). This was associated with a clear increase in total RNA polymerase II (Pol II) levels, as well as the transcriptionally active forms of Pol II, CTD-S5p and CTD-S2p (Fig. 3, C to H, fig. S9, A and B, fig. S11, B to D and fig. S8, D to F for PCA). Remarkably, Pol II recruitment at other chromosomes started to decay upon *CEN4\** excision (Fig. 3, D, F and H and fig. S11, B to D), suggesting that upregulation of gene expression at chromosome IV may titrate Pol II, and that Pol II might be present in limiting amounts in the cell. Finally, the characteristic wave-like spreading of unscheduled gene expression and histone acetylation from *CEN4\** to the arms of chromosome IV correlated with the spreading of H3K4me3, Pol II, Pol II CTD-S5p and CTD-S2p (fig. S12, B and C).

**Fig. 2.**
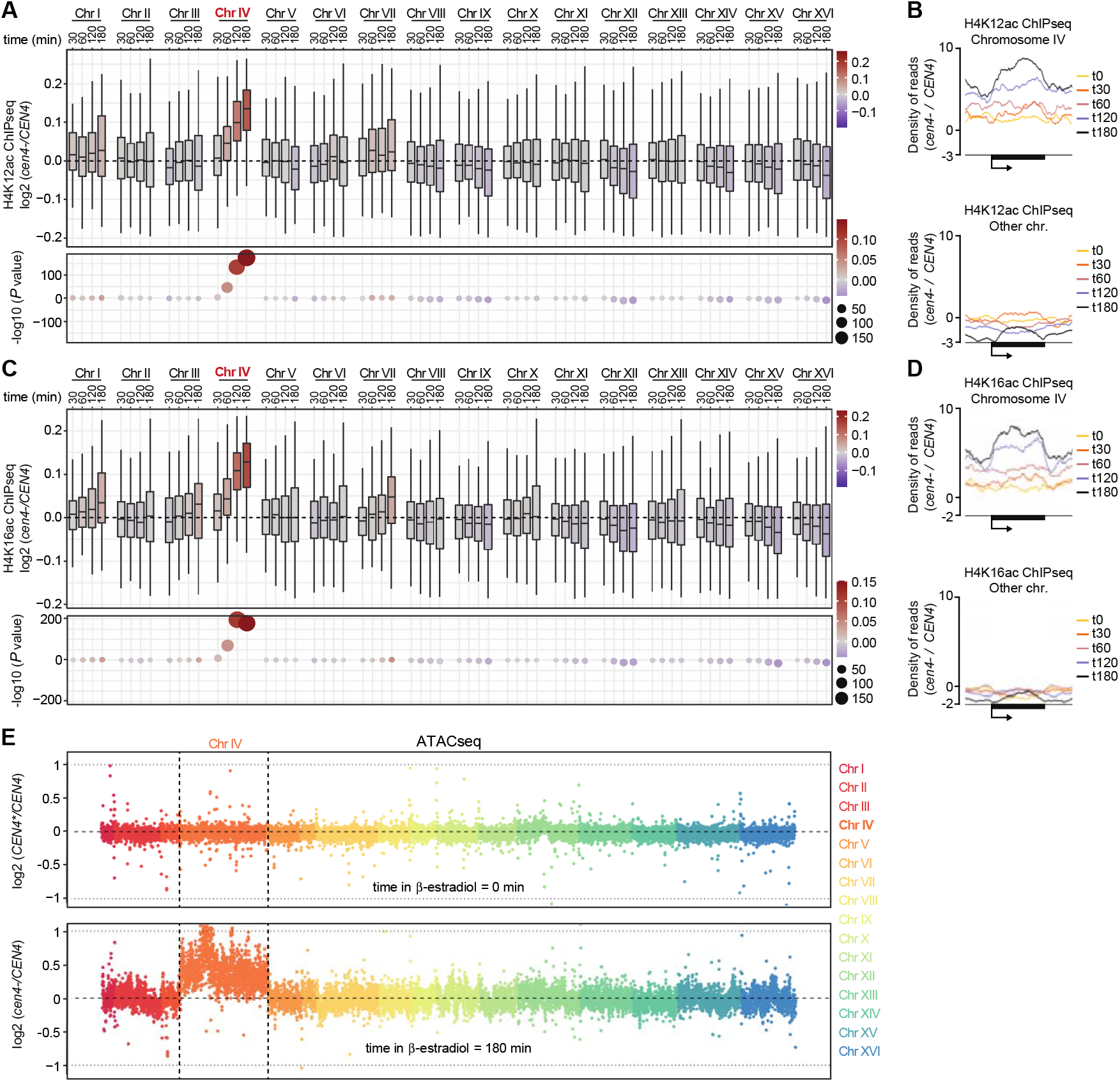
Centromere loss triggers chromatin relaxation in *cis*. H4K12ac **(A)** and H4K16ac **(C)** occurrence at each chromosome and at different time points after *CEN4\** excision. Metagene analysis of the top 20% ChIP signals showing increasing presence of H4K12ac **(B)** and H4K16ac **(D)** at chromosome IV genes and decreasing presence at genes on other chromosomes after centromere excision (see fig. S10 for metagnes of 100% of ChIP signals). **(E)** ATACseq experiments showing chromatin accessibility at time points 0 min (upper panel) and 180 min (lower panel) at each chromosome after *CEN4\** excision in *CEN4\** and *CEN4* cells. Each chromosome is represented using a dedicated color. *y* axis: chromosome coordinates. Each dot corresponds to bin of 10 Kb of chromosomal DNA.

**Fig. 3.**
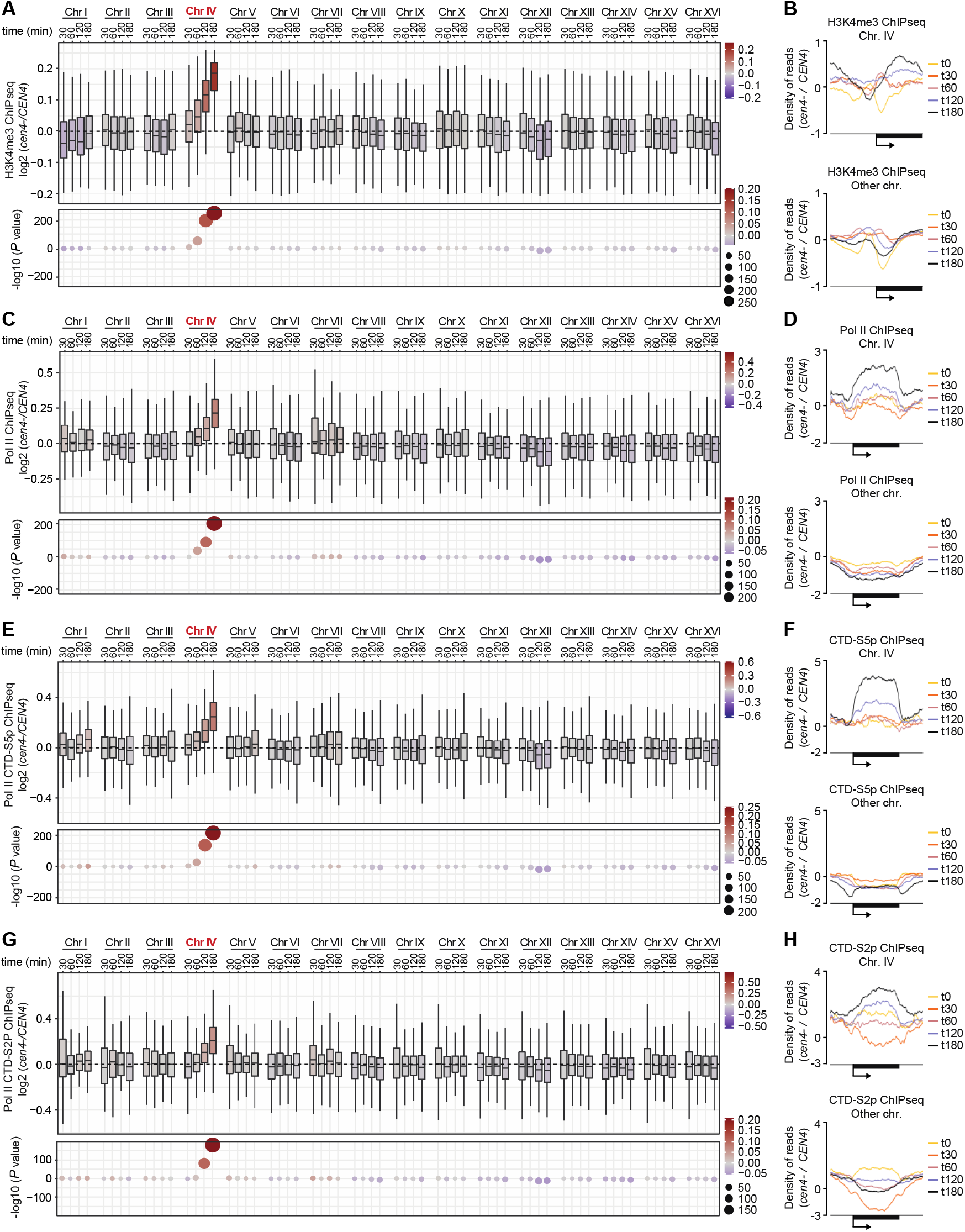
Lack of mitotic condensation triggers spontaneous transcriptional initiation. Assessment of H3K4me3 **(A)** Pol II **(C)** Pol II CTD-S5p **(E)** and Pol II CTD-S2p **(E)** occurrence using ChIPseq at each chromosome and at different time points after *CEN4\** excision. Metagene analysis of the top 20% ChIP signals shows increasing presence of H3K4me3 **(B)** Pol II **(D)** Pol II CTD-S5p **(F)** and Pol II CTD-S2p **(H)** at chromosome IV genes and a decrease at other chromosomes after centromere excision.

Supporting the idea that gene deregulation was a consequence of the condensation defect, unbiased hierarchical clustering analysis on RNAseq and ChIPseq data identified a cluster of genes at which an increase in positive histone marks and active Pol II clearly correlated with increased gene expression. This cluster was strongly and highly significantly enriched for chromosome IV genes (fig. S13A). Whereas some of the other clusters were enriched for certain GO terms, this “chromosome IV” cluster was not enriched for any particular pathway (fig. S13B). Furthermore, acetylation of histones on chromosome IV peaked 120 min after *CEN4\** excision (Fig. 2, A and B) while the occurrence of H3K4me3, Pol II, Pol II CTD-S5p and CTD-S2p was maximal at 180 min (Fig. 3, A, C, E and G). Thus, the failure to recruit Hst2 and condense the chromosome results first in increased histone acetylation and chromatin opening, and only subsequently in increased H3K4 methylation at promoters and transcription initiation, in the following interphase (fig. S13C).

Together, our data demonstrate that chromosome condensation has a key impact on the proper regulation of gene expression in the following interphase. Failure of a mitotic chromosome to condense in mitosis results in spontaneous recruitment of the transcription machinery, triggering unscheduled post-anaphase transcription of the entire chromosome, and unleashes genes from their control by upstream regulatory pathways (fig S14). Thereby, mitotic condensation resets gene expression at mitotic entry to safeguard against unlicensed transcription in the following interphase. It is interesting that a single locus, the centromere, instructs whole-chromosome gene silencing *in cis*. Whether centromeres only contribute to silencing via spreading of histone deacetylation or if they support other regulatory mechanisms like histone methylation remains to be investigated (*8, 9, 20–22*).

The effect of chromosome condensation on epigenetic markers and on gene expression in the ensuing interphase could be particularly relevant for asymmetrically dividing cells, such as stem cells, in which gene expression programs need to be reset to allow for maintenance of pluripotency, cellular identity and for determining cell fate (*4, 7, 20, 23–25*). Our study reveals an unexpected mechanism by which cells prevent post-mitotic transcriptional drifting, providing new inroads for exploration of mechanisms that maintain cellular homeostasis and control cell identity. Our results may also contribute to a better understanding of the etiology of diseases involving malfunctioning centromeres (*26, 27*).

## Acknowledgments

Sequencing was performed by the GenomEast platform, a member of the “France Genomique” consortium ANR-10-INBS-0009. We thank Dr. Ignacio Garcia for biostatistical insights.

## Funding

Norwegian Health Authority South-East grants 2017064, 2018012, 2019096 (JME, PC)

The Norwegian Cancer Society grants 182524, 208012 (JME)

Research Council of Norway grants 261936, 294916, 301268, 314811, 287911 (JME, PC,

MB)

Research Council of Norway Centers of Excellence funding scheme 262652 (JME)

French National Research Agency ANR-10-INBS-0009 (SLG, BJ)

Swiss National Science Foundation grant 31003-A-105904 (YB)

## Author contributions

Conceptualization: PC

Data curation: LRA, PH, SLG, XZ

Funding acquisition: MB, JME, PC

Formal analysis: PH, LRA, SLG, YB, PC

Methodology: BJ, YB, PC

Investigation: LRA, PC

Project administration: PC

Resources: MB, JME, YB, PC

Visualization: LRA, PH, XZ, PC

Validation: LRA

Supervision: BJ, PC, JME

Writing – original draft: LRA, PC

Writing – review & editing: LRA, PH, SLG, YB, JME, PC

## Data and materials availability

The RNAseq, ATACseq and H4K12ac, H4K16ac, H3K4me3, Pol II, Pol II CTD-S2p, Pol II CTD-S5p ChIPseq data have been deposited to the Gene Expression Omnibus with accession number GSE20225.

## Supplementary Materials

Materials and Methods

Figs. S1 to S14

Tables S1 to S5

References (*28*–*44*)

Captions of Data S1 to S3

