## Supplementary materials for "Mitotic chromosome condensation resets chromatin to maintain transcriptional homeostasis"

**This PDF file includes:**

Materials and Methods

Figs. S1 to S14

Tables S1 to S5

References (*28*–*44*)

Captions of Data S1 to S3

Materials and Methods

Strains and culture conditions

All strains used in this study (table S1) are from the S288c genetic background and were grown at 30°C in YPD medium [1% (w/v) yeast extract, 2% (w/v) peptone, 2% (w/v) dextrose].

The centromere excision assay

The strains used in this study express the Cre recombinase fused to the estradiol-binding domain, which retains Cre in the cytoplasm. Addition of 1 μM of β-estradiol (E8875, Sigma) to exponentially growing cells triggers nuclear import and thus activation of the Cre recombinase. Cells used in the different assays described in this study were collected at different time points after addition of β-estradiol.

The centromere of chromosome IV (*CEN4*) is flanked by lox recombination sites and is referred to as *CEN4** before excision and *cen4-* after excision by the Cre recombinase (Fig. 1A) (*1*). We use a control strain in which *CEN4* is not flaked with lox site to eliminate potential off target effects of β-estradiol during data analysis (Fig. 1A).

The same cre/lox system was used to excise a non-centromeric locus (*mCh**) from chromosome IV (fig. S5A).

*CEN4** and *mCh** excision efficiency assays

*CEN4** and *mCh** excision efficiency were assessed as in (*1*) with minor changes. Strains expressing the Cre recombinase fused to the estradiol-binding domain were treated with estradiol (E8875, Sigma) for different times before isolation of genomic DNA (gDNA) as following. Cell walls were degraded using Zymolyase (10 mg/ml; 120491-1, AMS biotechnology) in a cell resuspension buffer [50 mM KH2PO4 (pH 7.5), 1.2 M sorbitol] for 30 min at 30$^{\circ}$C, before addition of lysis buffer [100 mM Tris (pH 8.0), 50 mM EDTA, 1% SDS v/v]. Proteins were then precipitated using sodium acetate (1.1 M) and centrifuged at 14000 rpm. The supernatant was transferred to a new tube and the gDNA was precipitated using isopropanol. After centrifugation at 14000 rpm, pelleted gDNA was washed using 70% ethanol. After centrifugation at 14000 rpm, the pelleted gDNA was dried and resuspended in TE buffer [10mMTris (pH 8.0), 1mM EDTA] containing RNase DNase free (04716728001, Merck), and incubated overnight at room temperature. gDNA samples were analyzed by qPCR using a StepOnePlus™ Real-Time PCR system (Thermo Fisher Scientific), the Power SYBR Green PCR Master Mix (Thermo Fisher Scientific) and specific oligonucleotides (table S2). Data from three independent experiments were analyzed using the Applied Biosystems® Real-Time PCR Software (Life Technologies). *CLB6* has been used as a control gene (fig. S1A and fig. S5B). Results are presented as mean ± SD. Statistical significance was calculated using unpaired, two tailed Student’s t tests using GraphPad Prism (v7).

RNA-sequencing

Cells were collected from three independent experiments, snap frozen and stored at -80°C. Total RNA purification and genomic DNA removal were performed using the RNeasy Mini Kit (74104, Qiagen) and RNase-Free DNase Set (79254, QIAGEN), respectively, following manufacturer’s instructions. Total RNA quantification and quality control were performed using Tape Station 4150 (Agilent).

Library preparation and sequencing were performed at Novogene and raw data analysis at GeneVia. The quality of the RNAseq reads was inspected using the FastQC software [https://www.bioinformatics.babraham.ac.uk/projects/fastqc/] and multiQC software (*2*). RNAseq reads were aligned to the reference genome Ensembl *Saccharomyces cerevisiae* release R64-1-1 using STAR aligner version 2.6.1 (*3*). Gene-level read counts were obtained simultaneously with the alignment process using the corresponding *Saccharomyces cerevisiae* gtf annotation file from Ensembl.

Analysis of RNAseq data

Mapped RNA sequencing reads were imported into R and any reads with <1 mean reads over all samples were excluded. Pair-wise comparisons of 3 replicates for each strain at several time points were performed through the limma-voom pipeline (*4*). For each transcript at each time point, counts were first transformed by the limma::voom function and then fitted by limma::lmFit and limma::contrasts.fit. Empirical Bayes statistics for pairwise comparisons were generated by limma::eBayes. Differentially expressed genes (DEGs) as well as log2 fold change values were extracted by limma::topTreat, with a given *P* value parameter as threshold for DEGs (Data S1 and S2). Adjustment for multiple testing was in all cases done by setting the “adjust.method” to Benjamini-Hochberg (BH).

The samples were inspected using principal component analysis (PCA) of the log2-transformed normalised values (fig. S1B) and the results were visualised using ggplot2 R package (*5*).

S-phase arrest and release

To arrest cells in S phase, 200 mM of hydroxyurea (H8627, Sigma) was added to the culture medium (Fig. 1D). After 150 minutes, S-phase arrest was confirmed by visual inspection using a bright field microscope (cell doublets) (“S” in Fig. 1D). Then, β-estradiol was added and cultures were incubated during 90 minutes (“S+β-estradiol” in Fig. 1D). Cells were released from S-phase by washing 5x with YPD and then incubated for another 90 minutes (“Release” in Fig. 1D). In addition to visual inspection, an aliquot of cells was collected for cell cycle analysis by flow cytometry analysis (see below and fig. S7F). Cells were also collected at each time point, snap frozen and stored at -80°C for later RNA purification, reverse transcription and RT-qPCR experiments (Fig. 1E and fig. S7E).

G1-phase and M-phase arrest

To arrest cells in G1 phase, alpha-factor mating pheromone (RP01002, GenScript) has been added to a final concentration of 2 μg/ml (fig. S7A). After 60 minutes, the same amount of alpha-factor was re-added for another 60 minutes. G1-arrest was confirmed by visual inspection using a bright field microscope (cells are small, unbudded and shmoo formation can be observed) (“G1” in fig. S7A). Then, β-estradiol was added and cultures were incubated during 90 minutes (“G1+β-estradiol” in fig. S7A). An aliquot of cells has been collected for flow cytometry analysis at each time point (see below and fig. S7B). Cells were also collected at each time point, snap frozen and stored at -80°C for later RNA purification, reverse transcription and RT-qPCR experiments (fig. S7A).

To arrest cells in prometaphase of M phase, nocodazole (M1404, Sigma) has been added to a final concentration of 200 μg/ml together with β-estradiol (fig. S7C). After 180 minutes, M phase arrest was confirmed by visual inspection using a bright field microscope (cell doublets) (“M+β-estradiol” in fig. S7C). An aliquot of cells has been collected for flow cytometry analysis at each time point (see below and fig. S7D). Cells were also collected at each time point, snap frozen and stored at -80°C for later RNA purification, reverse transcription and RT-qPCR experiments (fig. S7C).

Cell cycle analysis by flow cytometry

Cell cycle profiling was performed using 10^7^ cells. After a wash step in water, cells were resuspended in 700 μl of water and fixed by slowly adding 700 μl of ethanol while vortexing. After 1 hour of incubation at room temperature, cells were resuspended in filtered 50 mM sodium citrate buffer pH 7.0 containing 250 µg/ml RNAse A and 1 mg/ml proteinase K and incubated overnight at 37°C. Cells were then resuspended in filtered 50 mM of sodium citrate [pH 7.0] and sonicated 3 seconds (30% output) in a bioruptor pico sonicator (Diagenode). Sytox green (S7020, Invitrogen) was then added to a final concentration of 1 µM. After 1 hour at room temperature, samples were analyzed by flow cytometry (488 nm excitation and emission peak of 523nm). Results were analyzed as previously published (*6*) using the BD LSRFortessa Cell Analyzer (BD Biosciences) and FlowJo Software (BD Biosciences) (fig. S7B, D and F).

Real-time quantitative PCR (RT-qPCR)

Total RNA purification was done using the RNeasy Mini Kit (74104, Qiagen) and the QuantiTect Reverse Transcription Kit (205311, Qiagen) following manufacturer’s instructions. RT-qPCRs were done using a StepOnePlus™ Real-Time PCR System (Thermo Fisher Scientific) and analyzed using the Applied Biosystems® Real-Time PCR Software (Life Technologies). A complete list of primers used for RT-qPCR can be found in table S2. The *MNN2* gene was used as a control gene because, based on the RNAseq data, its expression is not affected by the *CEN4** excision. Results are presented as mean ± SD. Experiments were performed at least 3 times and statistical significance was calculated using unpaired, two tailed Student’s t tests using GraphPad Prism (v7). *P* values < 0.05 were considered as significant.

Chromatin immunoprecipitation (ChIP)

Our ChIP procedure was adapted from (*1*). Cells were fixed with 1% (vol/vol) formaldehyde for 30 min. Formaldehyde was quenched for 5 min by adding glycine to a final concentration of 125 mM. Cells were washed with cold Tris-buffered saline, resuspended in lysis buffer [50 mM HEPES-KOH (pH 7.5), 140 mM NaCl, 1 mM EDTA, 1% (vol/vol) Triton X-100, 0.1% (vol/vol) Na-deoxycholate, and protease inhibitor cocktail] and were sonicated twice for 15 min by performing alternating cycles of 30 s pulses followed by a 30 s cool-down period using a Diagenode Bioruptor Pico. After centrifugation, supernatants were immunoprecipitated at 4$^{\circ}$C using anti-H4K12ac, H4K16ac, H3K4me3, Pol II, Pol II CTD-S2p and Pol II CTD-S5p (table S3) and protein A and G Dynabeads (10001D and 10003D, Thermo Fisher Scientific). Beads were first washed with lysis buffer, followed by washing with lysis buffer containing 500 mM NaCl, then with wash buffer [10 mM Tris (pH 8.0), 250 mM LiCl, 0.5% (vol/vol) Nonidet P-40, 0.5% (wt/vol) Na-deoxycholate and 1 mM EDTA], and finally with TE buffer [10 mM Tris (pH 8.0) and 1 mM EDTA]. Elution was performed in 50 mM Tris (pH 8.0), 10 mM EDTA, 1% (wt/vol) SDS at 65$^{\circ}$C, and the cross-link was reversed 5 hours at 65$^{\circ}$C. After RNase DNase free (04716728001, Merck) and proteinase K treatment (03115828001, Merk), DNA fragments were purified using QIAquick PCR purification kit (28104, Qiagen). DNA fragments were used for ChIPseq experiments (see below).

Library preparation and sequencing of ChIP samples (ChIPseq)

ChIP samples were purified using SPRIselect beads (Beckman-Coulter, Villepinte, France) and quantified using the Qubit 4 fluorimeter (Thermo Fischer Scientific, Illkirch, France). ChIPseq libraries were prepared from 2 to 10ng of double-stranded purified DNA using the MicroPlex Library Preparation kit v3 (C05010001, Diagenode, Seraing, Belgium), according to manufacturer's instructions. In the first step, the DNA was repaired to get molecules with blunt ends. Then, stem-loop adaptors with blocked 5 prime ends were ligated to the 5 prime end of the genomic DNA, leaving a nick at the 3 prime end. The adaptors cannot ligate to each other and do not have single-strand tails, avoiding non-specific background. In the final step, the 3 prime ends of the genomic DNA were extended to complete library synthesis and Illumina compatible indexes were added through a PCR amplification (7 or 10 cycles). Amplified libraries were purified and size-selected using SPRIselect beads (Beckman Coulter) at a ratio of 1/1 (= volume PCR/volume Beads) to remove free primers and other reagents. The purified libraries were sequenced on Illumina Hiseq 4000 sequencer as Single-Read 50 base reads following Illumina’s instructions. Image analysis and base calling were performed using RTA 2.7.7 and bcl2fastq 2.17.1.14. Adapter dimer reads were removed using DimerRemover [https://sourceforge.net/projects/dimerremover].

Analysis of ChIPseq data

Sequenced reads were mapped to the *Saccharomyces cerevisiae* genome (sacCer3) using Bowtie 1.0.0 (*7*) with the following arguments: -m 1 --strata --best -y -S -l 40 -p 8. Peaks were called using MACS2 v2.1.1.20160309 on the pool of replicates versus their corresponding input. MACS2 was run with default parameters except for: “-g 12000000 –nomodel –extsize 200”. For conditions H3K4me3_wt_60 and H3K4me3_mut_180, the additional parameters “–to-large” was added to correct for the high number of reads in the IPs compared to the input. Peaks were annotated relative to genomic features using Homer v4.11.0 (*8*) (annotations were extracted from gtf file downloaded from ensembl v102). Then, the number of reads falling in the union of all peak sets was generated using BEDtools (*9*) v2.26.0.

ChIP read counts were imported into R. The mean of replicates was used to calculate the log2 fold change between the indicated samples. Principal component analysis (PCA) was performed in R by first applying a variance stabilizing transformation to the raw data through the DESeq2::vst function (*10*). The 500 most variable genes across all samples were then used to calculate principle components by the stats::prcomp function. The first and second principal component was plotted for each replicate (fig. S8).

BigWig files were generated starting from .sam files containing mapped sequencing reads. Samtools was applied with parameters "view -bS -q 20" to generate .bam files that were further applied to bedtools using "genomecov -bg" parameters to generate .bedgraph files (*9, 11*). Finally, bedGraphToBigWig was used to generate bigWig files. Bigwig files in normalized reads per million (RPM) was normalized to input with bigwigCompare (V3.5.1). The size of the compared bins was 200bp. The difference of peak intensity was further calculated by subtracting in each time point the *CEN4* values from *CEN4** samples (Fig. 2, B and D and Fig. 3, B, D, F and H). In the cases indicated (fig. S10 and fig. S11) *CEN4** and *CEN4* values were both shown. The normalized read density profiles were generated with Easeq (V1.11) for metagene analysis.

Library preparation and sequencing for ATACseq experiments

ATAC-seq experiments were performed in duplicates as previously described (*12*). Briefly, 1–5 million yeast cells were collected and washed with spheroplasting buffer (SB) [40 mM HEPES pH 7.5, 10 mM MgCl2, 1 M sorbitol]. Then, cells were resuspended with 190 μL SB and 10 μL of 10 mg/mL zymolyase in SB. After 30 minutes incubation at 30°C, spheroplasted cells were washed twice with 200 μL of SB. Then, the pellet was resuspended in 50 μL transposition mix (25 μLTagment DNA Buffer, 22.5 μL nuclease-free water, 2.5 μL Nextera Tagment DNA Enzyme 1) using the Tagment DNA TDE1 Enzyme and Buffer Kit (Illumina 20034197). The solution was incubated at 37°C during 30 minutes and purified using the DNA Clean & Concentrator™-5 Kit (Zymo Research) and eluting in 11 μL of nuclease-free water.

For the library amplification, each PCR reaction had 25 μL NEBNext Hi-Fidelity 2× PCR Master Mix (New England Biolabs M0541S), 7.5 μL of nuclease-free water, 6.25 μL Universal Primer Ad1 (10 μM), 6.25 μL barcoded reverse primer (Ad2.[1–16]) (10 μM) (table S2 and (*13*) for ATAC-seq primer sequences) and 5 μL purified transposed DNA. The PCR amplification was: 1 cycle: 72°C for 5 min; 1 cycle: 98°C for 30 s; 10 cycles [98°C for 10 s, 63°C for 30 s, 72°C for 1 min]; hold at 4°C $\infty$.

PCR products were purified and size-selected using SPRIselect beads (Beckman-Coulter, Villepinte, France) to remove free primers and other reagents. This step was repeated twice using a ratio (volume of beads/volume of PCR) equal to 1.4/1. ATAC-seq libraries were sequenced on an Illumina HiSeq 4000 sequencer as paired-end 100 base reads. Image analysis and base calling were performed using RTA version 2.7.7 and bcl2fastq version 2.20.0.422.

Analysis of ATACseq data

Reads were mapped to *Saccharomyces cerevisiae* genome (assembly sacCer3) using the Encode ATAC pipeline v2.0.2 that uses bowtie2 (*14*) v2.3.4.3 to align reads. Briefly, Bowtie2 is used to map reads to the reference genome with default parameters except for “-X2000.” Then, samtools view v1.9 (*11*) is used to remove reads (unmapped, mate unmapped, not primary alignment reads, reads failing platform, low quality reads (MAQ <30)) with the following parameters “samtools view -F 1804 -q 30.” Then, duplicated reads are marked with Picard MarkDuplicates v2.20.7 [“Picard Tools - By Broad Institute.” n.d. Accessed August 26, 2016. http://broadinstitute.github.io/picard/] and removed with samtools with the following parameter “-F 1804.” Peak calling was done using the Encode ATAC pipeline v2.0.2 using macs2 v2.2.4 with the following parameters “–shift 75 –extsize 150 –nomodel -B –SPMR –keep-dup all –call-summits”. Peak set retained are the overlap / optimal ones.

To generate genomic coverage plots, Deeptools multiBamSummary v3.5 (*15*) was used to compute the number of reads in non-overlapping bins of 1Kb all over the Saccharomyces cerevisiae genome (assembly sacCer3). Then, R scripts with R v4.1.1 and the library ggplot2 v3.3.5 were used to generate plots.

Hierarchical clustering, heatmaps

Heatmaps showing hierarchical clustering of data were generated by the R pheatmap library [https://CRAN.R-project.org/package=pheatmap]. Clusters were generated by the pheatmap::pheatmap function using clustering_method="ward.D2". In all cases, the data was separated into three, five or ten clusters and the clustering was manually reviewed to find which separation gave clusters of meaningful size. Five clusters were selected in all the performed analysis. Genes of each cluster were extracted and analyzed further for over-representation of gene ontology or chromosome enrichment (fig. S2 and fig. S13). Colors of the heatmaps were the mean log2 fold change of the transcripts of that cluster. A heatmap was also used without performing any hierarchical clustering to show the mean expression changes of all transcripts on different chromosomes, normalized to t=0 (fig. S1D).

Gene ontology (GO) and chromosome over-representation analysis

Groups of genes (DEGs or hierarchically clustered genes) were analyzed for relative over-representation of genes belonging to functional categories (GO, biological processes) or being on specific chromosomes. The chromosome over-representation was performed by the piano::runGSAhyper function (*16*) using a gene set created manually of all genes included in the analyzed RNAseq data and their chromosome position. Gene ontology was performed using the clusterProfiler::enrichGO function (*17*) with the org.Sc.sgd.db gene ontology database (fig. S3 and fig. S13). The statistical test performed was in both cases a hypergeometric test (equivalent to a one-tailed Fisher exact test). Benjamini-Hochberg *P* value adjustment was performed to correct for multiple testing and relatively enriched categories were shown for *P*-adj < 0.05.

Binned positions on Chr IV plots

Plots showing bins along chromosome IV were employed to show the distribution of DEGs along the length of the chromosome for the RNA seq data (Fig. 1C and fig. S6C) or the log2 fold change values for ChIPseq data (fig. S12 and Data S3). Plots were made with a loess smoothed line using the ggplot2::geom_smooth function with the parameter method=”loess”. Grey bands show the 95% confidence intervals.

Box plots, dot plots, outlier exclusion

Box plots were generated in R by the ggplot2::geom_boxplot function (Fig. 1B, Fig. 2A and C, Fig. 3A, C, E and G, fig. S1E and fig. S5C). Default settings were used: middle line showing the median, upper and lower hinges showing 75^th^ and 25^th^ quartiles, upper and lower whiskers ending at 1.5*inter-quartile-range beyond the hinges. In some plots, it is noted in the figure legends that outliers were excluded in the visualization, meaning that individual points beyond the whiskers were not shown to improve visibility of the data when there were outliers that force the axis to be strongly extended. In these cases with box plots, outliers were excluded by setting outlier.shape=NA. Dot plots that show RNAseq or ChIP data signal along the length of a chromosome were also set to be limited to an upper and lower quantile values to increase the visibility of the trends in the data (fig. 6A and B and fig. S9). In these cases, quantiles were calculated in R by stats::quantile and points above 0.99 or below 0.01 are excluded. Outliers were only removed in visualizations and had no influence on any statistical testing.

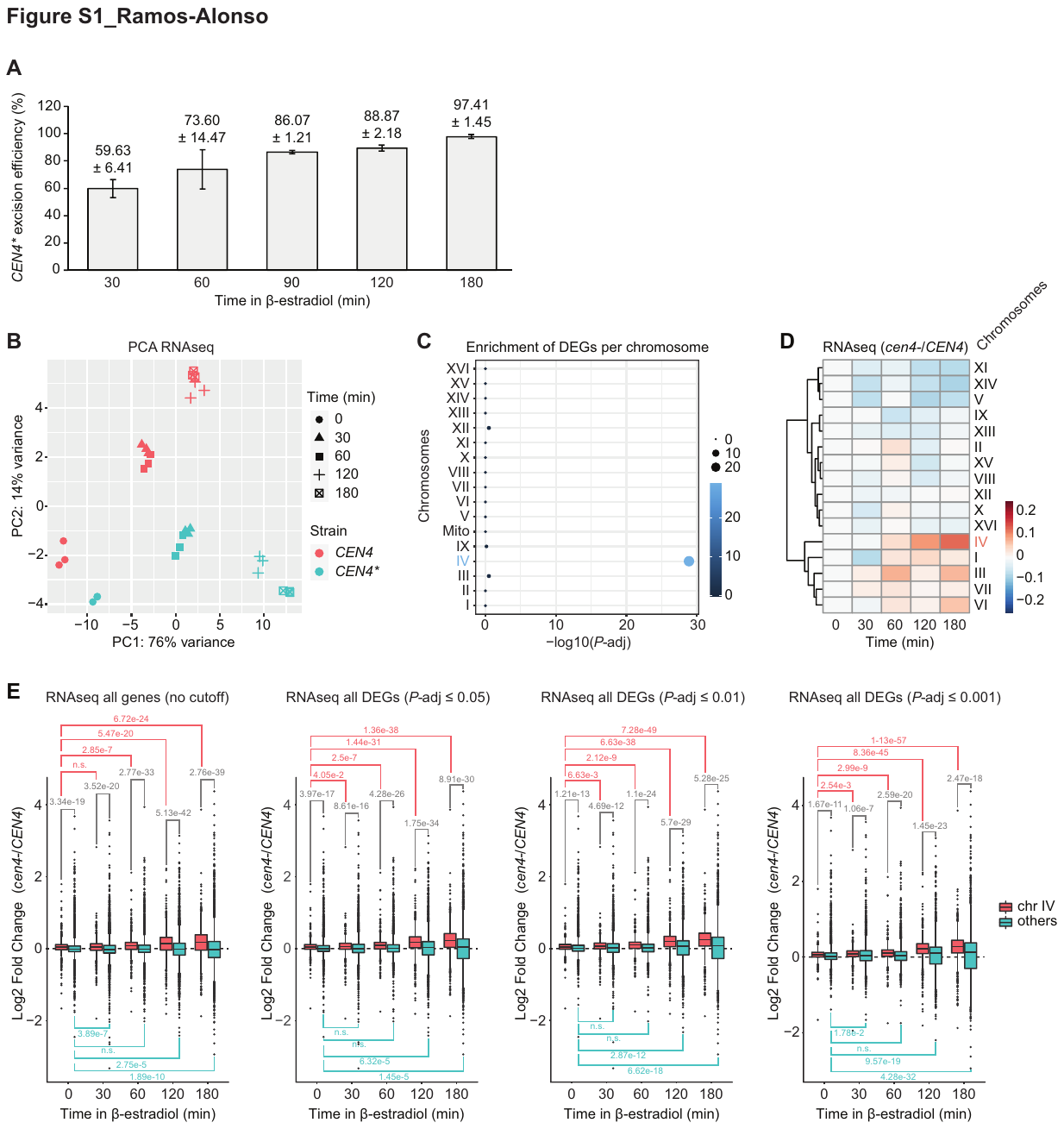

Fig. S1. *CEN4** excision correlates with a strong upregulation of gene expression in *cis*.

**(A)** Assessment of *CEN4** excision efficiency after 30, 60, 90, 120 and 180 minutes of the addition of β-estradiol. Error bars represent the SEM of three independent experiments (Mean (%) ± SEM)*.* **(B)** Principal component analysis (PCA) of RNAseq time-course experiment in *CEN4* and *CEN4** cells at 0, 30, 60, 120 and 180 minutes after the addition of β-estradiol. **(C)** Over-representation of differentially expressed genes (DEGs) at each chromosome compared to the total counts of genes on the chromosome. The size of the dot represents the –log10 of the *P* value in the *x* axis and it is also shown as color code. *P* values were calculated by a hypergeometric test and adjusted for multiple testing by the Benjamini–Hochberg method. DEGs were based on statistical differences between *cen4-* and *CEN4* at any time point by using a *P-*adj cutoff of 0.001. **(D)** Genome-wide heatmap of all transcripts from RNAseq time-course experiments showing the log2 fold change of *cen4-* normalized to *CEN4* (*cen4-/CEN4*) clustered by chromosome and normalized to time 0. Mean expression levels of the log2 fold change (*cen4-/CEN4*) of the transcripts in each cluster are indicated as color code. **(E)** Box plots of the RNAseq time-course experiment showing the log2 fold change (*cen4-/CEN4*) of the expression of the genes from Chr IV and the rest of chromosomes (Others). The different plots show (from left to right) all transcripts, or DEGs based on statistical differences between *cen4-* and *CEN4* in each time point by using a *P-*adj cutoff of 0.05, 0.01 and 0.001. *P* values calculated by Mann–Whitney U test are indicated for each comparison.

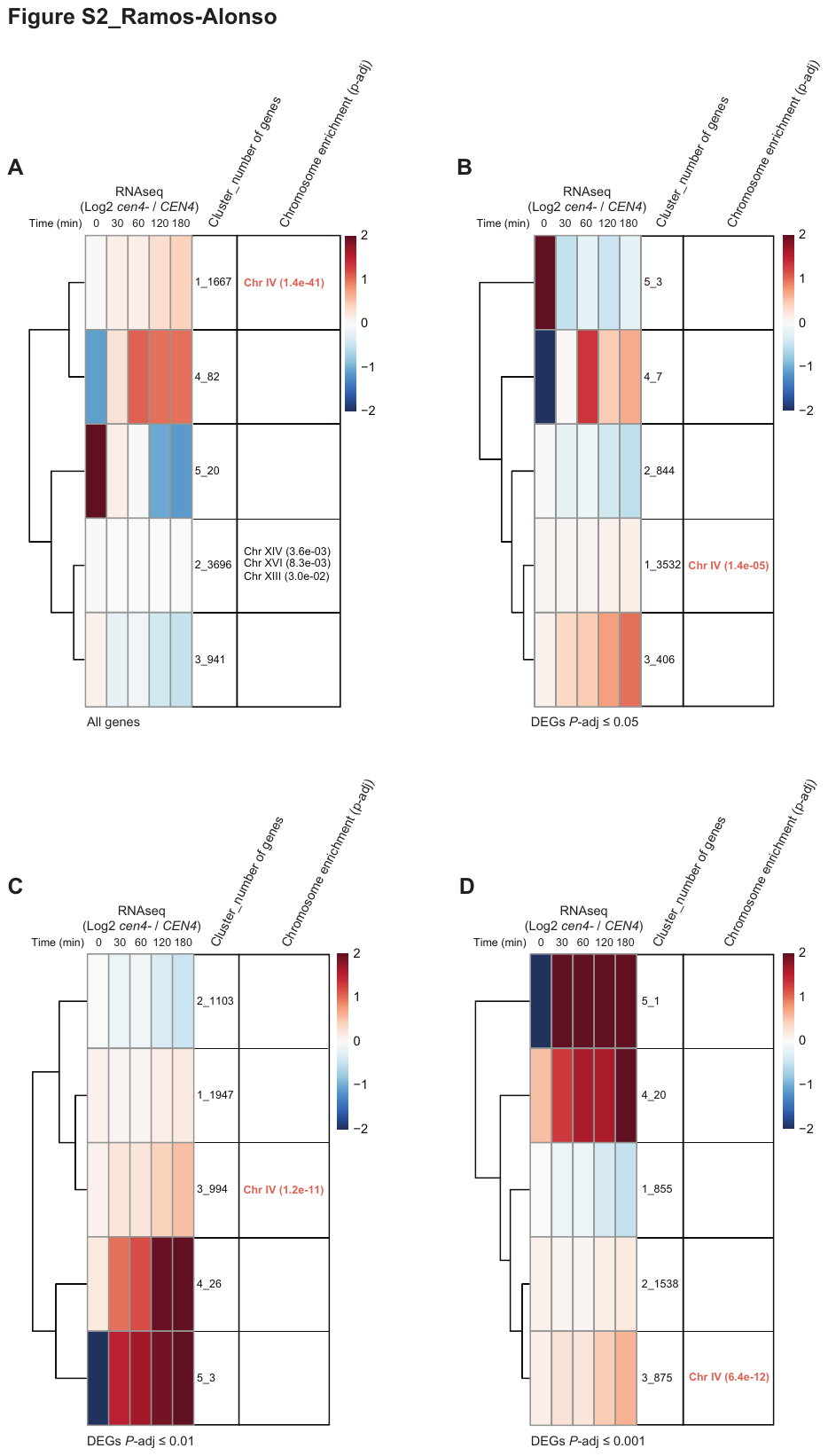

Fig. S2. Unbiased hierarchical clustering of RNAseq data. Genes with similar expression pattern in RNAseq time-course experiments are visualized using heatmaps. The log2 fold change (*cen4-/CEN4*) was calculated for each time point and mean expression levels in each cluster are indicated as a color code. The number of genes of each cluster are indicated next to the cluster number. Significant chromosome enrichment (*P*-adj <0.05) is indicated for each cluster. The *P* values were calculated by a hypergeometric test and adjusted for multiple testing by the Benjamini–Hochberg method. The different panels show all transcripts from RNAseq time-course experiment (A) or DEGs based on statistical differences between *cen4-* and *CEN4* in each time point by using a *P-*adj cutoff of 0.05 (B), 0.01 (C) and 0.001 (D).

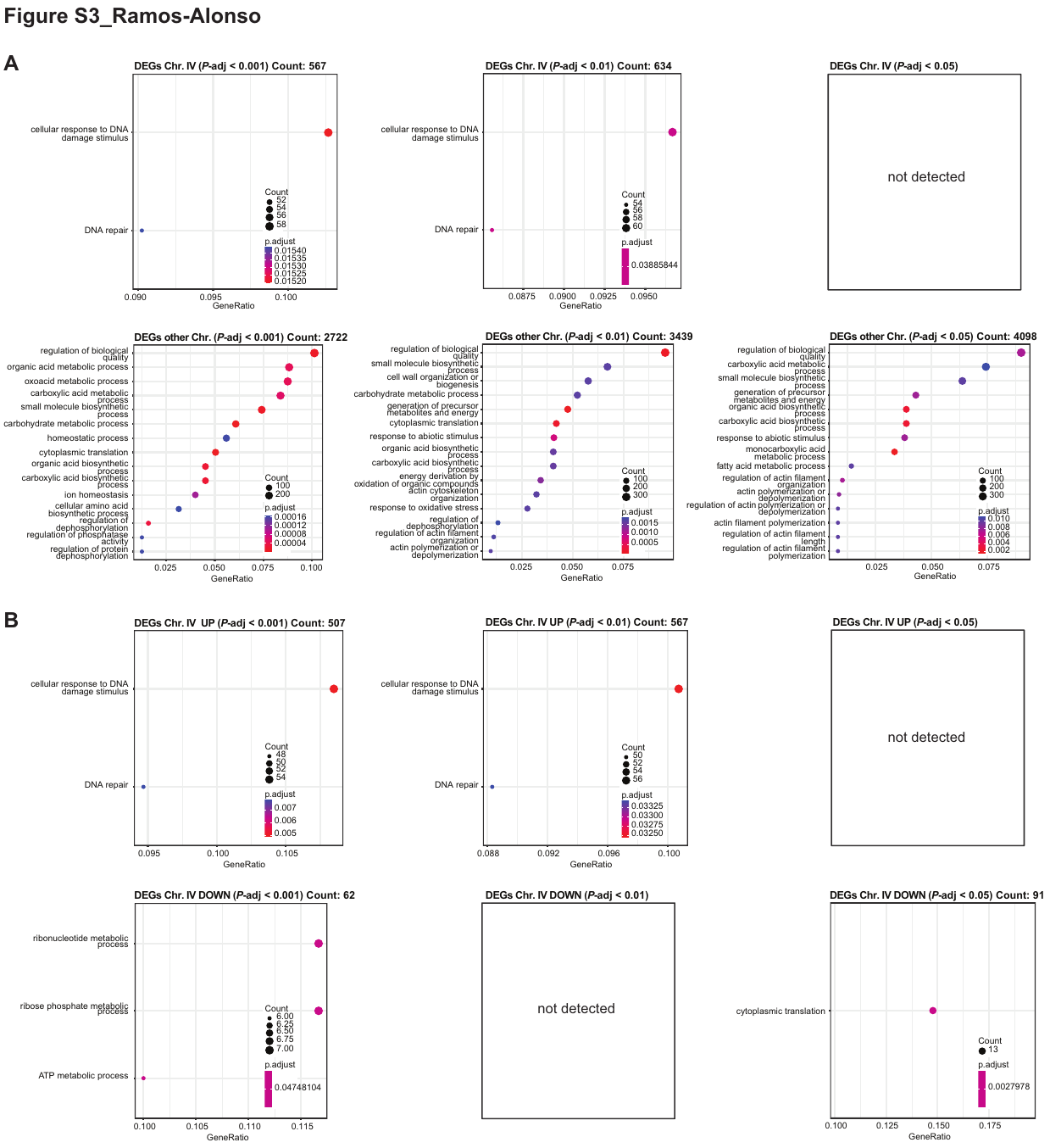

Fig. S3. Gene ontology (GO) analysis of RNAseq data. GO analysis showing the most significant GO-terms of (A) all genes from chromosome IV (upper panels) and DEGs from other chromosomes (lower panels) and (B) upregulated DEGs from chromosome IV (upper panels) and downregulated DEGs from chromosome IV (lower panels). In both panels, GO-terms are categorized by gene ratio (*x* axis), count of DEGs in each GO-term (size of the dot) and *P-*adj significance (color code). *P* values were calculated by a hypergeometric test and adjusted for multiple testing by the Benjamini–Hochberg method. DEGs were based on statistical differences between *cen4-* and *CEN4* at any time point by using a *P-*adj cutoff of 0.05, 0.01 and 0.001, in each case indicated.

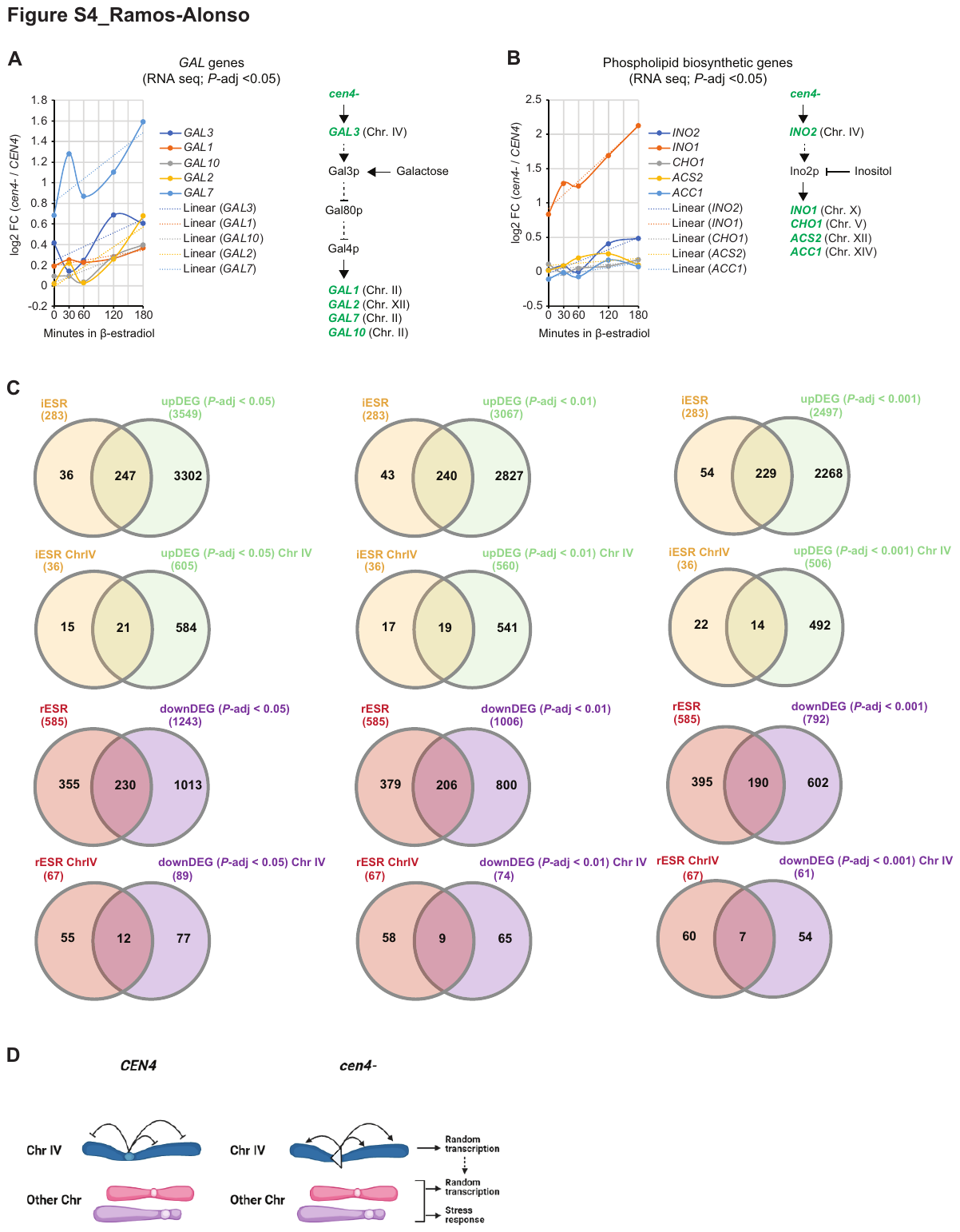

Fig. S4. *CEN4** excision triggers activation of regulatory pathways in absence of stimuli and correlates with a stress response-like gene expression response. (A) Left: *CEN4** excision triggers expression of *GAL* genes in absence of galactose. Right: The *GAL* genes pathway. (B) Left: *CEN4** excision triggers expression of inositol biosynthetic genes in while inositol levels are not affected in YPD medium. Right: The inositol genes pathway. In both panels (A) and (B) results are shown as log2 fold change of *cen4-* / *CEN4* (*P-*adj cutoff 0.05). Genes in green are upregulated upon *CEN4** excision. (C) Venn diagrams showing overlap of induced ESR genes (iESR, in yellow) and repressed ESR genes (rESR, in red) (*18*) and DEGs from RNAseq time-course experiments (upregulated in green, downregulated in purple). DEGs were isolated based on statistical differences between *cen4-* and *CEN4* at each time point by using a *P-*adj cutoff of 0.05, 0.01 and 0.001, as indicated. (D) Model showing that centromere excision triggers unscheduled and excessive transcription in *cis*, which may affect genes on other chromosomes, leading to activation of pathways in absence of cognate stimuli and possibly stress response.

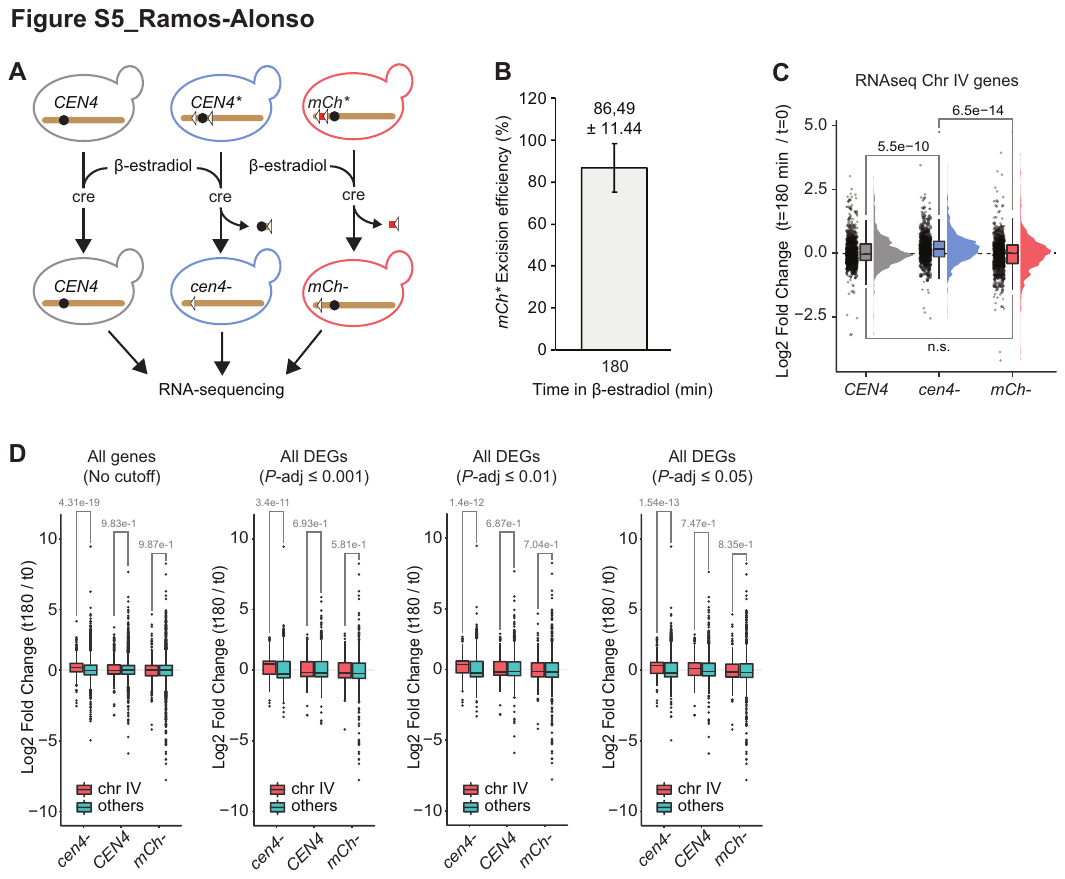

Fig. S5. Excision of a non-centromeric patch of DNA does not alter gene expression. (A) Experimental layout. Left: *CEN4* control strain. Middle: *CEN4** strain with loxed *CEN4*. Addition of β-estradiol triggers the excision of *CEN4** (*cen4-*). Right: *mCh** strain with loxed *mCherry.* Addition of β-estradiol triggers the excision of *mCh** (*mCh-*). (B) Calculation of *mCh** excision efficiency (Mean (%) ± SEM) after 180 minutes of the addition of β-estradiol. Error bars represent the SEM of three independent experiments. (C) Rainfall Plot of RNAseq showing the log2 fold change of the expression of the genes from Chr IV after 180 minutes of the addition of β-estradiol normalized to time 0 (t180/t0) in the strains indicated in (A). (D) Box plots of the same RNAseq experiment as in (C) showing the log2 fold change (t180/t0) of the expression of the transcripts from Chr IV and the rest of chromosomes (Others) using (from left to right): all transcripts (no cutoff), DEGs based on statistical differences between t180 and t0 by using a *P*-adj cutoff of 0.001, 0.01 or 0.05. In both (C) and (D). *P* values calculated by Mann–Whitney U test are indicated for each comparison.

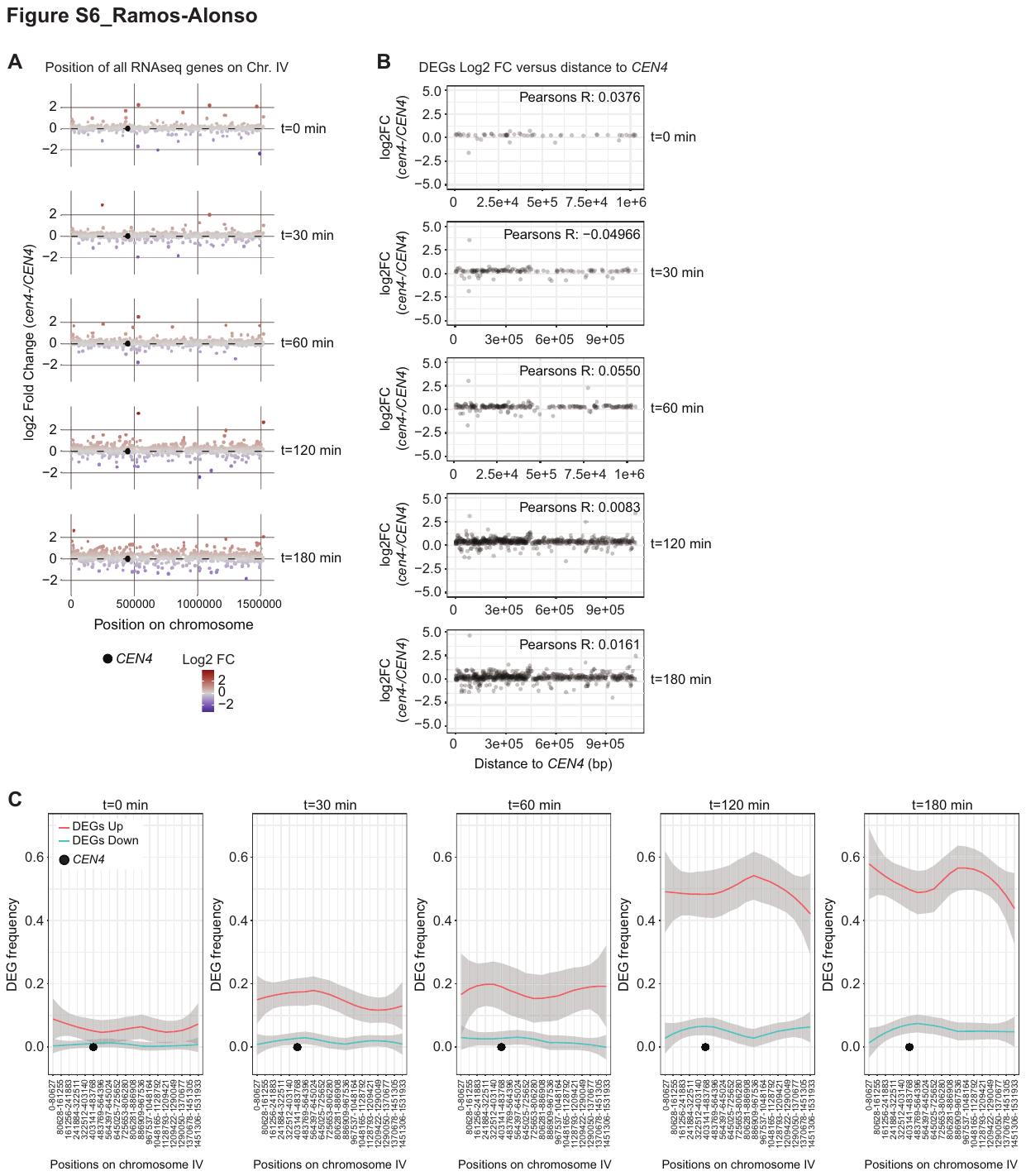

Fig. S6. *CEN4** excision spreads gene upregulation along chromosome arms. (A) Dot plots showing the log2 fold change (*cen4-/CEN4*) from all transcripts at chromosome IV (*y* axis) and their position on the chromosome (*x* axis) at each time point upon *CEN4** excision. The log2 fold change values are shown on the *y* axis and color coded. The highest (above 99.9% quantile) and lowest (below 0.1% quantile) extreme datapoints were excluded in the visualization to improve the visibility of the data. (B) Plots of the of the RNAseq time-course experiment showing the log2 fold change (*cen4-/CEN4*) from DEGs of chromosome IV (*y* axis) and absolute distance to the centromere of Chr IV (*x* axis) in each time point. Pearson correlation coefficients between log2 fold change (*cen4-/CEN4*) and distance to the centromere are indicated. (C) Representation of the frequency of chromosome IV DEGs upregulated (red) and downregulated (teal) in each time point of the RNAseq time-course experiment by different bins of positions on chromosome IV. In panels (B) and (C) DEGs were based on statistical differences between *cen4-* and *CEN4* at each time point by using a *P-*adj cutoff of 0.001.

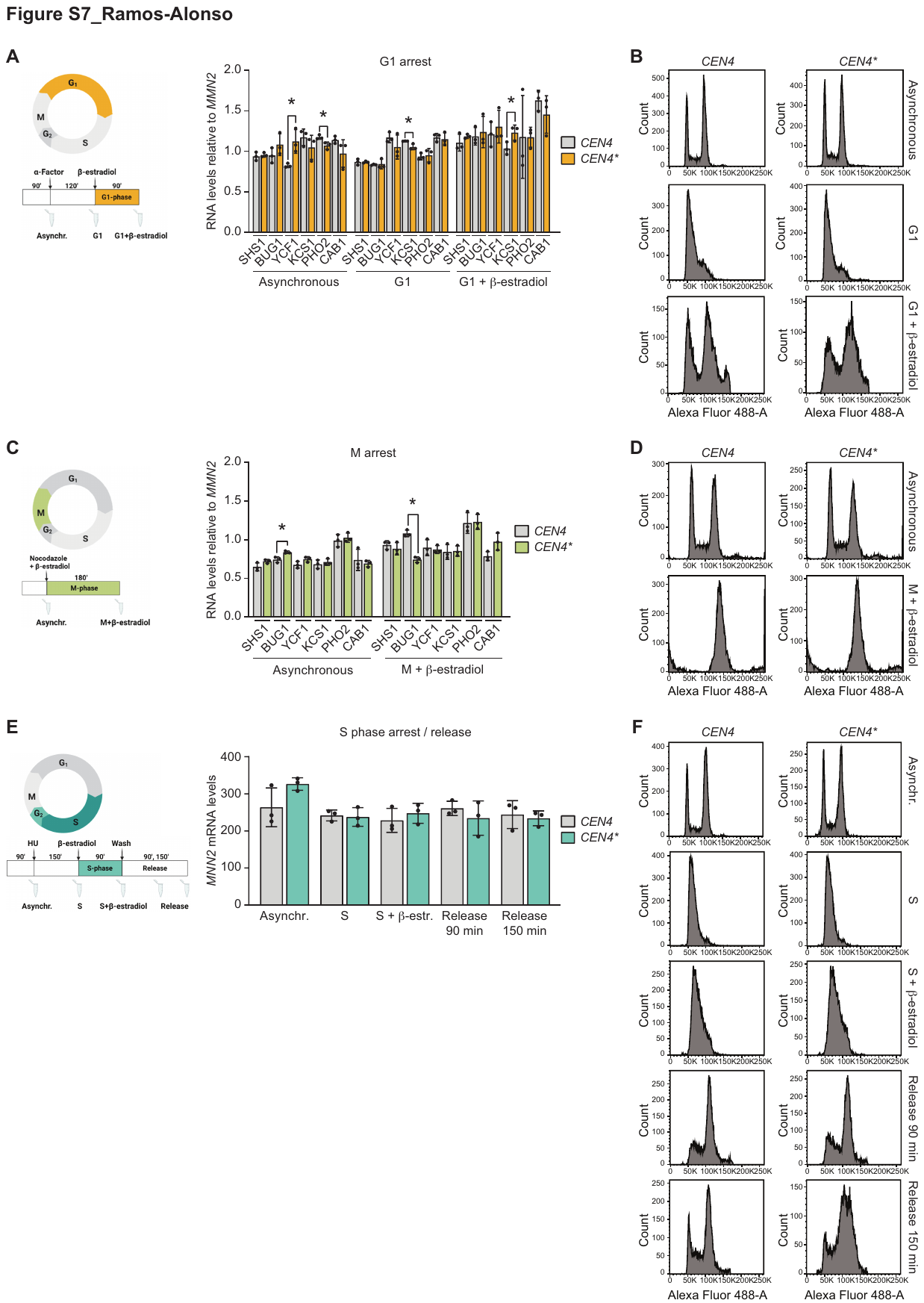

Fig. S7. Cell cycle arrest experiments. RT-qPCR analysis of the expression of six selected genes from chromosome IV at different time points: (A) during G1-phase arrest and (C) during prometaphase arrest. **P* values < 0.05 considered as significant. (E) Expression of the control gene *MNN2* during S-phase and release experiments was analyzed using RT-qPCR. (B), (D) and (F) show the analysis of the cell cycle using flow cytometry performed with the cells indicated in (A), (C) and (E), respectively.

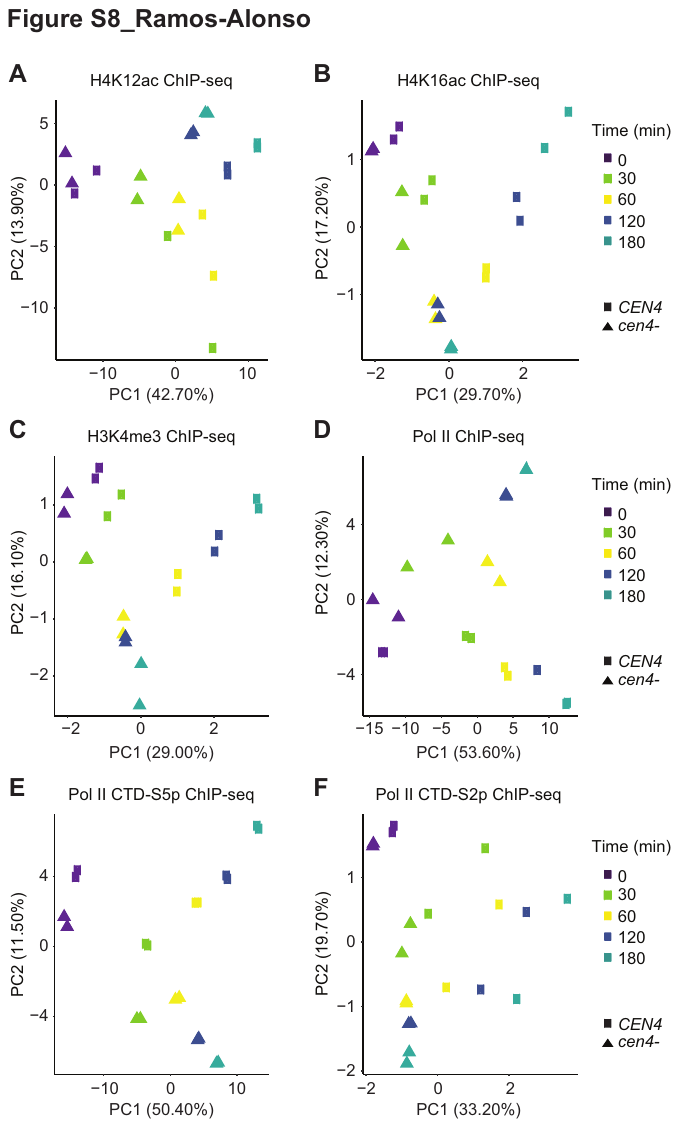

Fig. S8. Principal component analysis (PCA) of ChIPseq data. ChIPseq H4K12ac (A), H4K16ac (B), H3K4me3 (C), Pol II (D), Pol II CTD-S5p (E) and Pol II CTD-S2p (F). First, the union of all peak positions in all the genes has been performed, and then a computation of the number of reads falling in the peaks has been made by using BEDtools (*9*) v2.26.0. Finally, a PCA was computed on regularized logarithm transformed data calculated with the method proposed by (*10*).

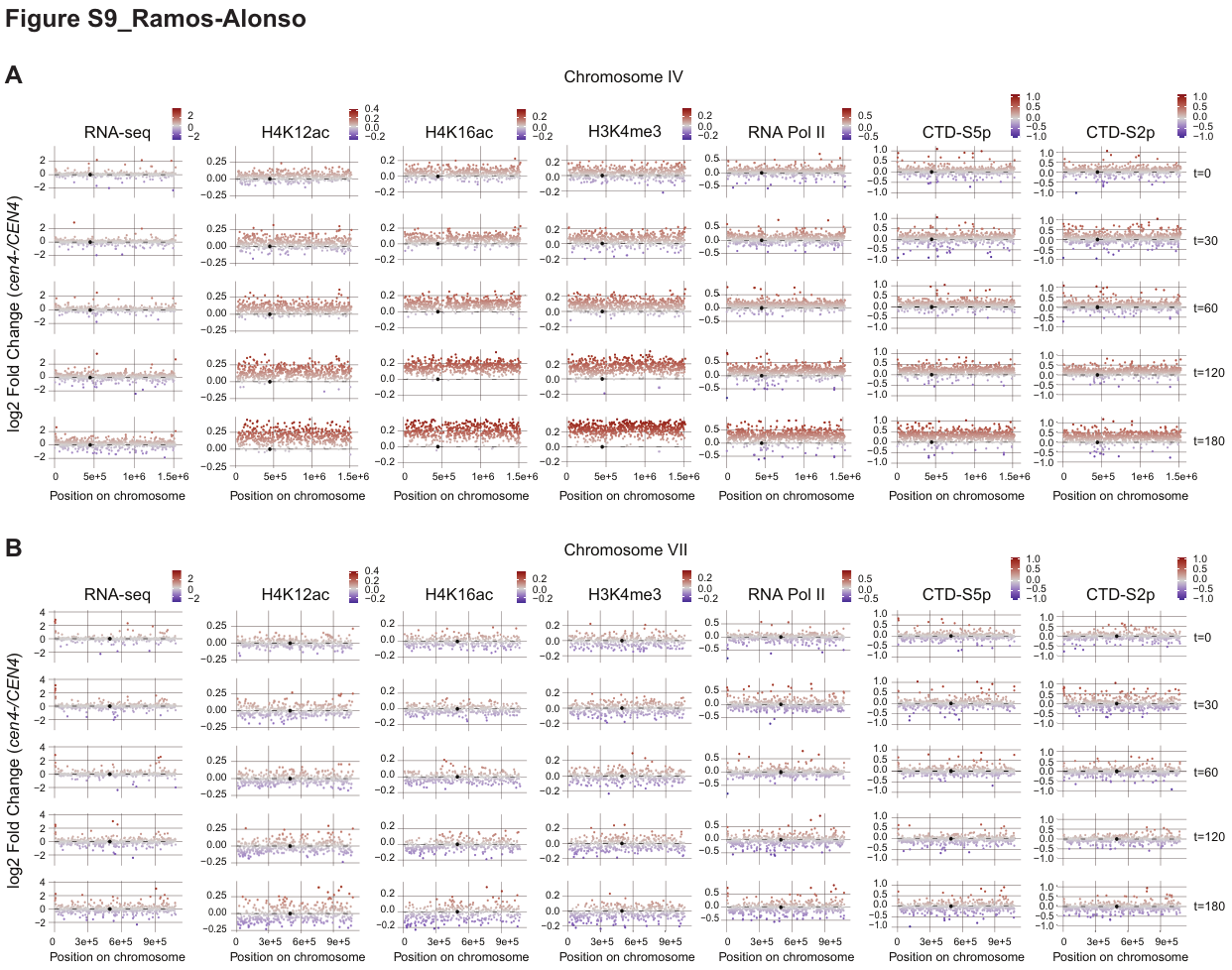

Fig. S9. Centromere excision elicits histone acetylation and transcriptional initiation in *cis*.

Dotplots of the RNAseq and ChIPseq time-course experiments showing the log2 fold change (*cen4-/CEN4*) from all genes at Chr IV **(A)** and Chr VII, an example of another chromosome of comparable size **(B)**. *x* axis: chromosome coordinates. *y* axis: log2 fold change (also indicated as color code). The highest (above 99.9% quantile) and lowest (below 0.1% quantile) extreme datapoints were excluded in the visualization to improve the visibility of the trends.

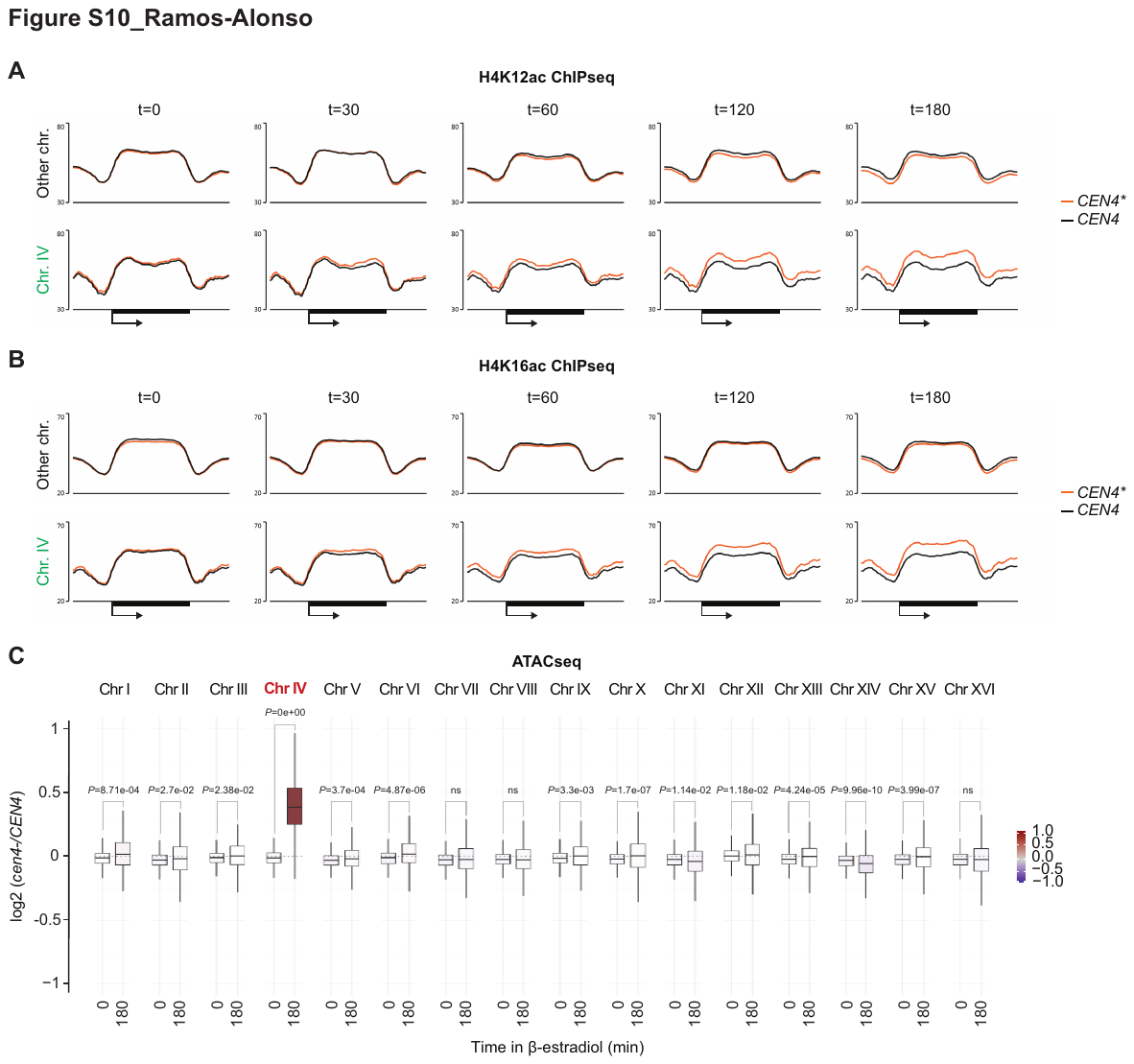

Fig. S10. Centromere excision elicits histone acetylation and chromatin opening in *cis*. The mean profiles of H4K12ac (A) and H4K16ac (B) were generated for all genes at other chromosomes (upper panels) and chromosome IV (lower panels) at each time point after *CEN4** excision in *CEN4** and *CEN4* cells. Black bar represents genes from TSS (arrow) to TTS. (C) ATACseq results presented as box plots showing the increase of chromatin accessibility at chromosome IV before (t = 0) and after (t=180 min) *CEN4** excision in *CEN4** and *CEN4* strains.

Fig. S11. Metagene analysis of H3K4me3, Pol II, CTD-S5p and CTD-S2p ChIPseq experiments. The mean profiles of H3K4me3 (A), Pol II (B), CTD-S5p (C) and CTD-S2p (D) were generated for all genes at other chromosomes (upper panels) and chromosome IV (lower panels) at each time point after *CEN4** excision in *CEN4** and *CEN4* cells. Black bar represents genes from TSS (arrow). In (A), ChIP signals are shown $\boldsymbol{-}$/$\boldsymbol{+}$ 1500 bp relative to TSS.

Fig. S12. Centromere excision spreads chromatin opening and transcriptional initiation along the chromosome arms. Representation of the relative occurrence of H4K12ac and H4K16ac (A), H3K4me3(B), and Pol II, CTD-S5p and CTD-S2p (C) at chromosome IV at different time points after *CEN4** excision. *x* axis: chromosome coordinates; *y* axis: log2 fold change (*cen4-/CEN4*).

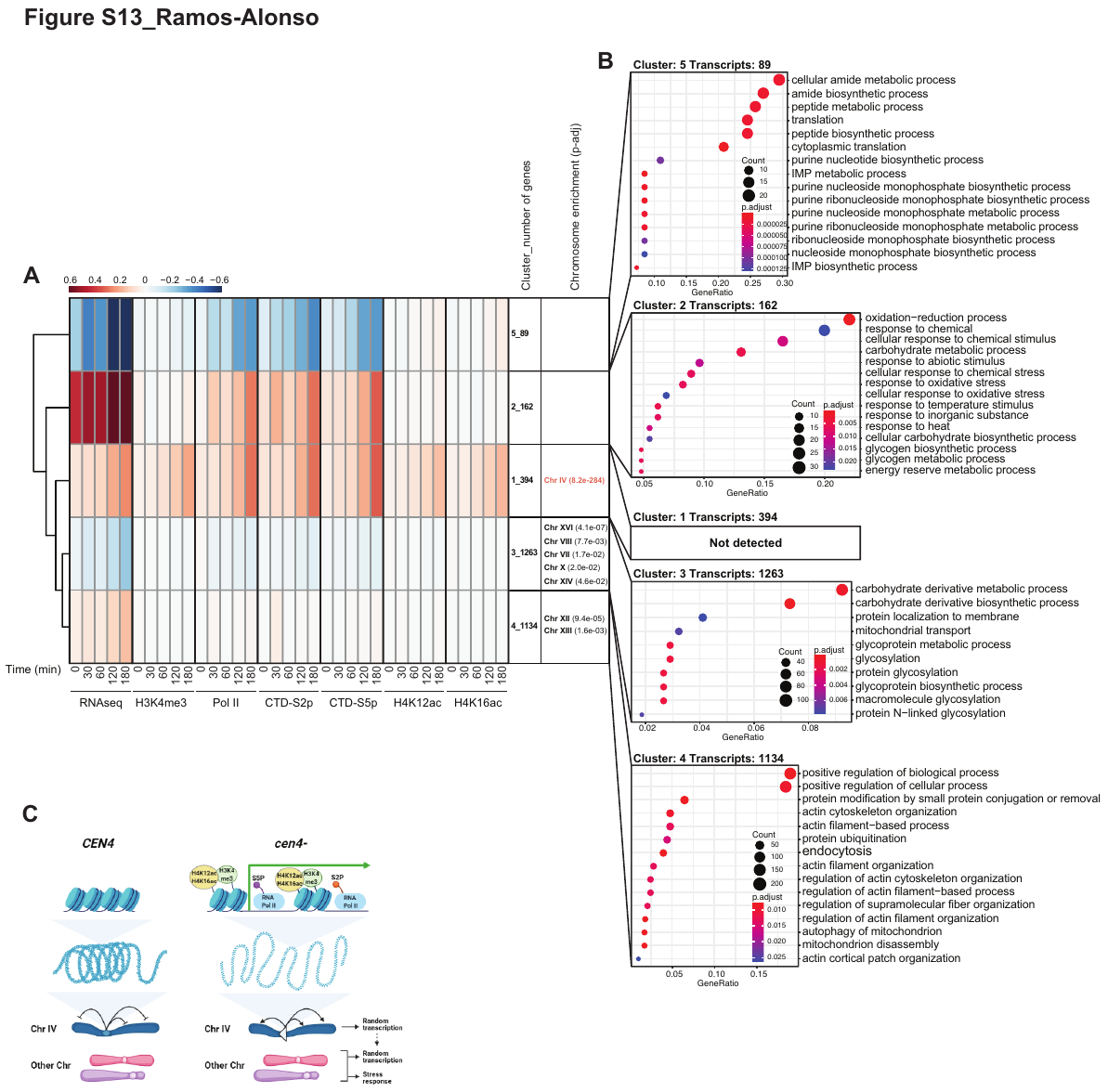

Fig. S13. Unbiased clustering of RNAseq and ChIPseq results. (A) Hierarchical clustering of RNAseq and ChIPseq time-course experiments. Mean expression [log2 fold change (*cen4-/CEN4*)] are indicated as a color code. The number of genes in each cluster are indicated next to the cluster number. Significant chromosome enrichment (*P*-adj <0.05) is indicated for each cluster. (B) GO analysis indicating the most significant terms in each cluster from (A). The GO-terms are categorized by gene ratio (*x* axis), count of DEGs in each GO-term (size of the dot) and *P-*adj significance (color code). For both the chromosome and GO enrichment, *P* values were calculated by a hypergeometric test and adjusted for multiple testing by the Benjamini–Hochberg method. (C) Failure to condense chromosome IV results in increased histone acetylation, which is followed by increased H3K4 methylation at promoters and initiation of transcription by RNA polymerase II.

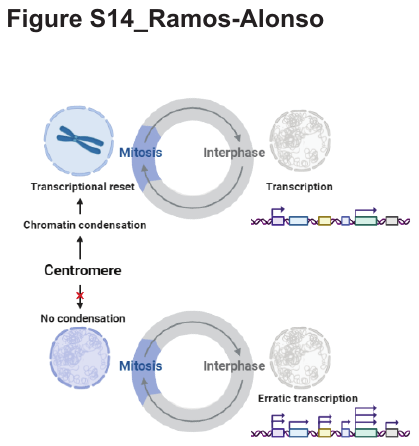

**Fig. S14. Centromere-induced mitotic chromosome condensation resets the transcriptome to prevent transcriptional drifting during interphase.**

| **Strain ID** | **Genotype** | **Source** |
| --- | --- | --- |
| yPC8 (*CEN4** or *cen4-*) | *MATa lox-CEN4-lox:KanMX GDP-creEBD78:LEU2 his3Δ1 leu2Δ0 ura3Δ0 met15Δ0* | YYB 9867 in (*1*) |
| yPC11 (*CEN4*) | *MATa ura3::GDP-cre EBD78:LEU2 his3Δ1 leu2Δ0 ura3Δ0 met15Δ0* | This study |
| yPC30 (*mCh** or *mCh-*) | *MATa trp1::TetO:TRP1 lys4::LacO:hphNT1 his3::LacR-GFP:HIS3 TetR-mCherry leu2::Cre-EBD78:LEU2 Shs1-loxPmCherry:Cln2(3’UTR)-NatNT2-loxP-GFP ade2* | yYB 14417 in (*1*) |

Table S1. Yeast strains (*S. cerevisiae*) used in this study

| **Oligonucleotide** | **Sequence (5’ – 3’)** | **Source** | **Use** |
| --- | --- | --- | --- |
| *MNN2*-F | GGCTACGGCTCTTCTATCT | This study | RT-qPCR |
| *MNN2*-R | CTTATCACCTTCACCAGCAG |  |  |
| *SHS1*-F | CCAGAGCGTACCAAGTTAAG |  |  |
| *SHS1*-R | ACTGGGCCTCTAACTCTTC |  |  |
| *BUG1*-F | CTGTCGATGACCTACAGGAT |  |  |
| *BUG1*-R | CTTGCTGCCTTAGCTTCTG |  |  |
| *KCS1*-F | CCTTGTGCACTGGATCTAAA |  |  |
| *KCS1*-R | GCCTAAACGCCTAGAAGTG |  |  |
| *PHO2*-F | GCTAACAGTCACCAGATCAC |  |  |
| *PHO2*-R | GCGTCATTTACCTGTCTACC |  |  |
| *CAB1*-F | CATGGACTCAAAGGCTATCTAC |  |  |
| *CAB1*-R | GAAGAACCGCCTACTCTACTA |  |  |
| P1b*-*F (*CEN4**) | CCAATTGGCTGGACGTAAAC |  |  |
| P2b*-*R (*CEN4**) | TTTGGCCGCTCCTAGTG |  |  |
| *CLB6-*F | GGAACATGGCCAAATTCATCATGG | (*1*) | *CEN4** and *mCh** excision efficiency |
| *CLB6-*R | GGCGATATACATGGACATTGCAGC |  |  |
| P148-R (*mCh**) | AATTGGGACAACTCCAGTGAAA | This study |  |
| P149-F (*mCh**) | GAGTTGATGAATCTCGGTGTGT |  |  |
| *Ad1_noMX* | AATGATACGGCGACCACCGAGATCTACACTCGTCGGCAGCGTCAGATGTG |  |  |
| *Ad2.1_TAAGGCGA* | CAAGCAGAAGACGGCATACGAGATTCGCCTTAGTCTCGTGGGCTCGGAGATGT |  |  |
| *Ad2.2_CGTACTAG* | CAAGCAGAAGACGGCATACGAGATCTAGTACGGTCTCGTGGGCTCGGAGATGT | (*13*)  (*13*) | ATACseq  ATACseq |
| *Ad2.3_AGGCAGAA* | CAAGCAGAAGACGGCATACGAGATTTCTGCCTGTCTCGTGGGCTCGGAGATGT |  |  |
| *Ad2.4_TCCTGAGC* | CAAGCAGAAGACGGCATACGAGATGCTCAGGAGTCTCGTGGGCTCGGAGATGT |  |  |
| *Ad2.5_GGACTCCT* | CAAGCAGAAGACGGCATACGAGATAGGAGTCCGTCTCGTGGGCTCGGAGATGT |  |  |
| *Ad2.6_TAGGCATG* | CAAGCAGAAGACGGCATACGAGATCATGCCTAGTCTCGTGGGCTCGGAGATGT |  |  |
| *Ad2.7_CTCTCTAC* | CAAGCAGAAGACGGCATACGAGATGTAGAGAGGTCTCGTGGGCTCGGAGATGT |  |  |
| *Ad2.8_CAGAGAGG* | CAAGCAGAAGACGGCATACGAGATCCTCTCTGGTCTCGTGGGCTCGGAGATGT |  |  |
| *Ad2.9_GCTACGCT* | CAAGCAGAAGACGGCATACGAGATAGCGTAGCGTCTCGTGGGCTCGGAGATGT |  |  |
| *Ad2.10_CGAGGCTG* | CAAGCAGAAGACGGCATACGAGATCAGCCTCGGTCTCGTGGGCTCGGAGATGT |  |  |
| *Ad2.11_AAGAGGCA* | CAAGCAGAAGACGGCATACGAGATTGCCTCTTGTCTCGTGGGCTCGGAGATGT |  |  |
| *Ad2.12_GTAGAGGA* | CAAGCAGAAGACGGCATACGAGATTCCTCTACGTCTCGTGGGCTCGGAGATGT |  |  |
| *Ad2.13_GTCGTGAT* | CAAGCAGAAGACGGCATACGAGATATCACGACGTCTCGTGGGCTCGGAGATGT |  |  |
| *Ad2.14_ACCACTGT* | CAAGCAGAAGACGGCATACGAGATACAGTGGTGTCTCGTGGGCTCGGAGATGT |  |  |
| *Ad2.15_TGGATCTG* | CAAGCAGAAGACGGCATACGAGATCAGATCCAGTCTCGTGGGCTCGGAGATGT |  |  |
| *Ad2.16_CCGTTTGT* | CAAGCAGAAGACGGCATACGAGATACAAACGGGTCTCGTGGGCTCGGAGATGT |  |  |

Table S2. Oligonucleotides used in this study.

| **Antibody** | **Description** | **µg of chromatin used** | **Source** | **Cat#** |
| --- | --- | --- | --- | --- |
| Anti-RNAPol II | Anti-RNA Polymerase II Antibody, CTD Antibody, clone 8WG16, mouse monoclonal | 250 | Sigma-Aldrich | 05-952-I |
| Anti-CTD-S5p | RNA pol II CTD phospho Ser5 antibody (mAb), rat monoclonal | 100 | Active Motif | 61085 |
| Anti-CTD-S2p | RNA pol II CTD phospho Ser2 antibody (mAb), rat monoclonal |  |  | 61083 |
| Anti-H3K4me3 | Anti-Histone H3 (tri methyl K4) antibody - ChIP Grade (ab8580), rabbit polyclonal | 50 | Abcam | ab8580 |
| Anti-H4K12ac | Anti-acetyl-Histone H4 (Lys12) Antibody, rabbit polyclonal |  | Sigma-Aldrich | 07-595 |
| Anti-H4K16ac | Histone H4K16ac antibody (pAb), rabbit polyclonal |  | Active Motif | 39167 |

**Table S3. Antibodies used in this study.**

| RNAseq | *P*-adj Cutoff | Genes Chr IV | Genes Other Chr | Total genes | % Genes Chr IV | % Genes Other Chr | % Total DEGs | % Up Genes | % Down Genes | % Up Genes Chr IV | % Down Genes Chr IV |
| --- | --- | --- | --- | --- | --- | --- | --- | --- | --- | --- | --- |
| t0 | None | 810 | 5596 | 6406 | 12.6 | 87.4 | N/A | 69.5 | 30.5 | 81.4 | 18.6 |
| t30 |  |  |  |  |  |  |  | 65.6 | 34.4 | 80.0 | 20.0 |
| t60 |  |  |  |  |  |  |  | 68.2 | 31.8 | 81.5 | 18.5 |
| t120 |  |  |  |  |  |  |  | 72.0 | 28.0 | 83.9 | 16.3 |
| t180 |  |  |  |  |  |  |  | 68.2 | 31.8 | 82.7 | 17.3 |
| t0 | 0.05 | 205 | 897 | 1102 | 18.6 | 81.4 | 17.2 | 79.1 | 20.9 | 93.2 | 6.8 |
| t30 |  | 360 | 1660 | 2020 | 17.8 | 82.2 | 31.5 | 75.2 | 24.8 | 88.3 | 11.7 |
| t60 |  | 388 | 1463 | 1851 | 21.0 | 79.0 | 28.9 | 76.0 | 24.0 | 91.0 | 9.0 |
| t120 |  | 602 | 3125 | 3727 | 16.2 | 83.8 | 58.2 | 78.2 | 21.8 | 89.4 | 10.6 |
| t180 |  | 644 | 3511 | 4155 | 15.5 | 84.5 | 64.9 | 74.0 | 26.0 | 88.2 | 11.8 |
| t0 | 0.01 | 108 | 535 | 643 | 16.8 | 83.2 | 10.0 | 76.4 | 23.6 | 93.5 | 6.5 |
| t30 |  | 246 | 1111 | 1357 | 18.1 | 81.9 | 21.2 | 75.0 | 25.0 | 89.4 | 10.6 |
| t60 |  | 278 | 953 | 1231 | 22.6 | 77.4 | 19.2 | 74.8 | 25.2 | 92.1 | 7.9 |
| t120 |  | 541 | 2548 | 3089 | 17.5 | 82.5 | 48.2 | 79.5 | 20.5 | 90.6 | 9.4 |
| t180 |  | 582 | 3001 | 3583 | 16.2 | 83.8 | 55.9 | 75.4 | 24.6 | 90.0 | 10.0 |
| t0 | 0.001 | 58 | 322 | 380 | 15.3 | 84.7 | 5.9 | 78.7 | 21.3 | 89.7 | 10.3 |
| t30 |  | 148 | 716 | 864 | 17.1 | 82.9 | 13.5 | 74.8 | 25.2 | 89.2 | 10.8 |
| t60 |  | 177 | 605 | 782 | 22.6 | 77.4 | 12.2 | 73.0 | 27.0 | 90.4 | 9.6 |
| t120 |  | 482 | 1932 | 2414 | 20.0 | 80.0 | 37.7 | 80.2 | 19.8 | 90.5 | 9.5 |
| t180 |  | 520 | 2449 | 2969 | 17.5 | 82.5 | 46.3 | 75.7 | 24.3 | 90.8 | 9.2 |

Table S4. RNAseq DEGs at all time points after *CEN4** excision. DEGs were isolated based on statistical differences between *cen4-* and *CEN4* at each time point using a *P-*adj cutoff of 0.001, 0.01 or 0.05. Data are available for DEGs at chromosome IV and at other chromosomes.

| *P*-adj Cutoff | DEGs Chr IV | DEGs Other Chr | Total DEGs | % DEGs Chr IV | % DEGs Other Chr | % Total DEGs | % Up DEGs | % Down DEGs | % Up DEGs Chr IV | % Down DEGs Chr IV |
| --- | --- | --- | --- | --- | --- | --- | --- | --- | --- | --- |
| 0.05 | 694 | 4098 | 4792 | 14.5 | 85.5 | 74.8 | 74.1 | 25.9 | 87.2 | 12.8 |
| 0.01 | 634 | 3439 | 4073 | 15.6 | 84.4 | 63.6 | 75.3 | 24.7 | 88.3 | 11.7 |
| 0.001 | 567 | 2722 | 3289 | 17.2 | 82.8 | 51.3 | 75.9 | 24.1 | 89.2 | 10.8 |

Table S5. Sum of all RNAseq DEGs after *CEN4** excision. DEGs were isolated based on statistical differences between *cen4-* and *CEN4* at each time point using a *P-*adj cutoff of 0.001, 0.01 or 0.05. Data are available for DEGs at chromosome IV and at other chromosomes.

Data S1. (Separate file)

**The log2 fold change (*cen4-/CEN4*) of RNAseq time-course experiment.** The file contains the log2 fold change (*cen4-/CEN4*) and the *P-*adj of *cen4-/CEN4* comparison. For the expression of all genes in each time point (allGenes_t0; allGenes_t30; allGenes_t60; allGenes_t120; allGenes_t180). For DEGs based on statistical differences between *cen4-* and *CEN4* at each time point by using a *P-*adj cutoff of 0.05 (DEGs(0.05)_t0; DEGs(0.05)_t30; DEGs(0.05)_t60; DEGs(0.05)_t120; DEGs(0.05)_t180), DEGs by using a *P-*adj cutoff of 0.01 (DEGs(0.01)_t0; DEGs(0.01)_t30; DEGs(0.01)_t60; DEGs(0.01)_t120; DEGs(0.01)_t180) and DEGs by using a *P-*adj cutoff of 0.001 (DEGs(0.001)_t0; DEGs(0.001)_t30; DEGs(0.001)_t60; DEGs(0.001)_t120; DEGs(0.001)_t180). And for DEGs based on statistical differences between *cen4-* and *CEN4* at any time point by using a *P-*adj cutoff of 0.05 (DEGs(0.05)_any_timepoint), 0.01 (DEGs(0.01)_any_timepoint) and 0.001 (DEGs(0.001)_any_timepoint) including the mean value of all time points.

Data S2. (Separate file)

**The log2 fold change (t180/t0) of RNAseq control experiment**. The file contains the log2 fold change (t180/t0) and the *P-*adj of t180/t0 comparison for *CEN4*, *cen4-* and *mCh-*.

Data S3. (Separate file)

**The log2 fold change (*cen4-/CEN4*) of ChIPseq time-course experiment.** The file contains the log2 fold change (*cen4-/CEN4*) in each time point for Pol II, CTD-S5p, CTD-S2p, H3K4me3, H4K12ac and H4K16ac ChIPseq datasets.

5.	H. Wickham, C. Sievert, SpringerLink, in *Use R!* (Springer

Springer International Publishing : Imprint: Springer, Cham, 2016).
